# Only a fraction of UCP1 is required to sustain adaptive nonshivering thermogenesis in the cold

**DOI:** 10.64898/2026.08.06.743367

**Authors:** Qimuge Naren, Celso Pereira Batista Sousa-Filho, Weijun Pang, Natasa Petrovic

## Abstract

To address the long-standing question of the respective physiological contributions of classical brown *versus* beige adipocytes to adaptive nonshivering thermogenesis, we generated mice with lineage-specific ablation of UCP1 in thermogenic adipocytes of myogenic origin. This selectively targeted the major classical brown adipocyte lineage while preserving UCP1 expression in the remaining thermogenic adipocytes, reducing total UCP1 content by approximately 80 %. Unexpectedly, despite this profound reduction in UCP1 abundance, cold acclimation-recruited thermogenic capacity, assessed by adrenergic stimulation, remained largely preserved. In contrast, complete UCP1 deficiency abolished the adrenergically induced thermogenic response, demonstrating that UCP1 is indispensable for adaptive nonshivering thermogenesis. These findings indicate that in cold-acclimated mice only a fraction of the UCP1 normally present is required to sustain maximal thermogenic capacity. We further establish that the capacity to support UCP1-dependent oxidative metabolism, rather than UCP1 abundance, is the principal constraint on maximal thermogenic output under these conditions.

## Introduction

Although the vast majority of total UCP1 resides in classical brown rather than beige adipocytes^1,2^, the striking recruitment of beige adipocytes during cold exposure (e.g.^3–8^) has led to the widespread assumption that these adipocytes are major contributors to adaptive nonshivering thermogenesis. However, the relative physiological contributions of these two thermogenic adipocyte populations have not been directly determined. Because classical brown and beige adipocytes coexist within and across different adipose depots^9^, depot-based approaches such as surgical removal or denervation cannot distinguish their cell type-specific contributions to adaptive nonshivering thermogenesis^10,11^.

Genetic ablation of UCP1^12^ has established the indispensable role of UCP1 in adaptive thermogenesis^13,14^ but this approach cannot distinguish the physiological contributions of classical brown and beige adipocytes because UCP1 is eliminated from all thermogenic adipocytes. A diphtheria toxin receptor-based strategy has also been attempted^15^. Importantly, the distinct developmental origins of classical brown and beige adipocytes (myogenic and non-myogenic, respectively)^9,16–18^ enable a lineage-specific genetic approach to selectively disrupt UCP1-dependent thermogenesis in the thermogenic adipocytes of myogenic origin, which comprise the majority of classical brown adipocytes. Using this model, we determined the physiological consequences of eliminating UCP1 from ‘myogenic’ adipocytes while preserving UCP1 function in the remaining ‘non-myogenic’ thermogenic adipocytes. Unexpectedly, despite reducing total UCP1 content by nearly 80 %, cold acclimation-recruited adrenergically induced nonshivering thermogenic capacity remained largely preserved. In contrast, as expected, adaptive nonshivering thermogenesis was essentially abolished in mice lacking UCP1 globally. These findings together demonstrate that although UCP1 is indispensable for adaptive nonshivering thermogenesis, in cold-acclimated mice only a fraction of the UCP1 normally present is required to sustain maximal thermogenic capacity. Instead, maximal thermogenic output is constrained by the capacity of thermogenic adipose tissue to support oxidative metabolism.

## Results

### Myf5-Cre–mediated UCP1 deletion abolishes UCP1 protein expression in the majority of brown adipocytes within IBAT

To address the long-standing question of the contributions of classical brown *versus* beige adipocytes to adaptive nonshivering thermogenesis, we created mice in which UCP1 was selectively ablated in the thermogenic adipocytes of myogenic origin, which comprise the majority of classical brown but not beige adipocytes^9,16^. We crossed mice carrying floxed *Ucp1* alleles (Ucp1^fl/fl^) with a Cre-driver line expressing Cre under the control of the Myf5 promoter (Myf5-Cre), generating UCP1^Myf5cKO^ mice (Myf5-Cre^+/-^; Ucp1^fl/fl^) (Figure 1a). Details of the generation of the conditional *Ucp1* allele are provided in Figure S1.

**Figure 1.**
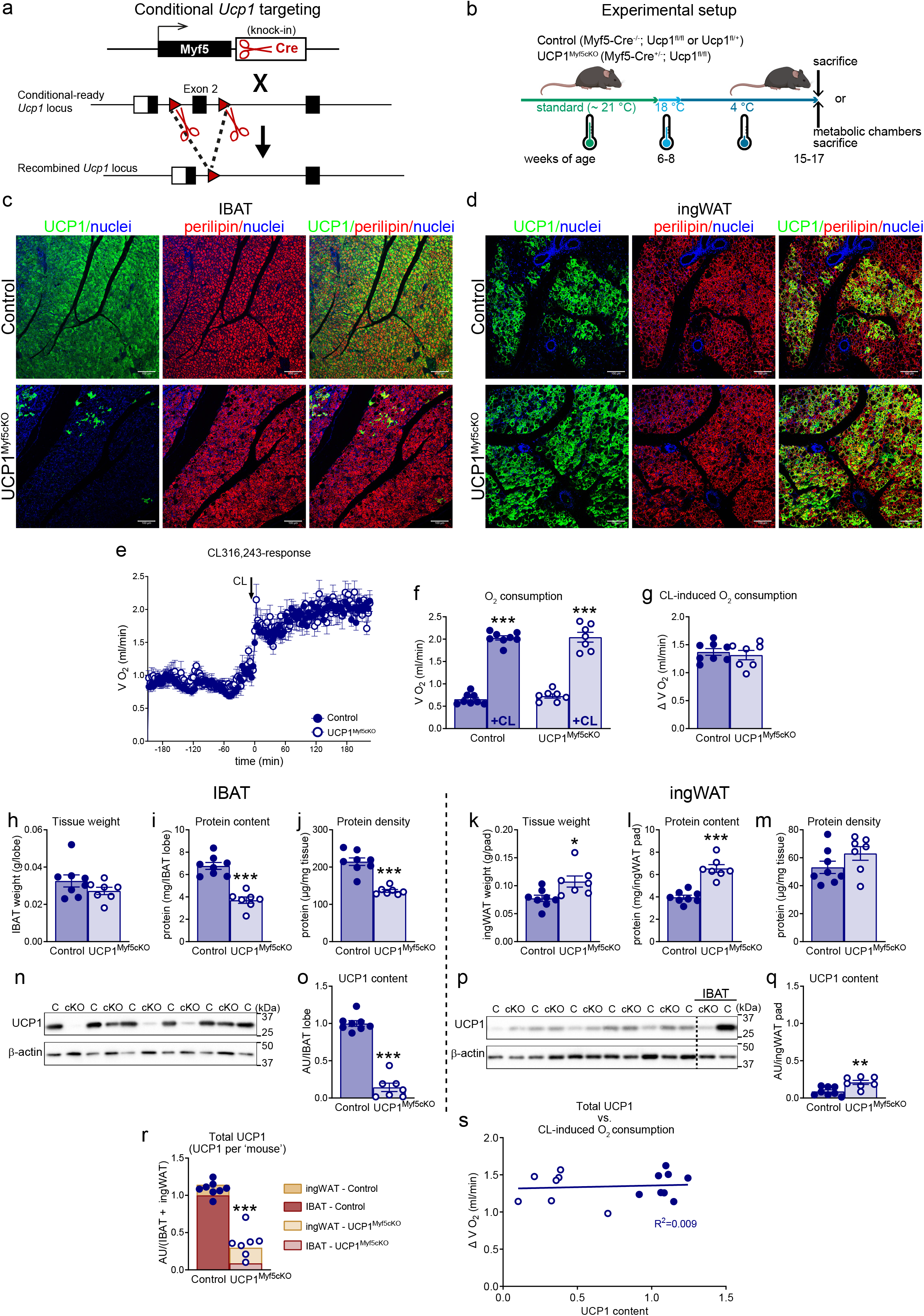
Preserved thermogenic capacity despite profound reduction in UCP1 content in cold-acclimated UCP1^Myf5cKO^ mice. **a,** Genetic strategy used to generate UCP1^Myf5cKO^ mice. **b,** Experimental design. **c,d,** Representative confocal images of interscapular brown adipose tissue (IBAT; **c**) and inguinal white adipose tissue (ingWAT; **d**) from control (upper panels) and UCP1^Myf5cKO^ mice (lower panels) acclimated to 4 °C. Sections were stained for UCP1 (green), perilipin (red), and nuclei (blue). Scale bar, 100 μm. **e–g,** Thermogenic capacity assessed by indirect calorimetry following CL316,243 (CL) administration. Male and female mice were analyzed together because no sex-dependent differences were observed (Figure S2c,d). Control (n = 8) and UCP1^Myf5cKO^ (n = 7) mice were housed overnight at 30 °C in the Promethion indirect calorimetry system, injected with CL (1 mg/kg), and monitored for a further 4 h. **e,** Oxygen consumption before and after CL injection. **f,** Baseline oxygen consumption (left bars) and mean oxygen consumption during the final 2 h after CL injection (right bars). **g,** CL-induced increase in oxygen consumption calculated as the difference between post- and pre-injection values shown in f. **h–j,** IBAT wet weight (**h**), total protein content (**i**), and protein density (**j**). **k–m,** Corresponding analyses of ingWAT wet weight (**k**), total protein content (**l**), and protein density (**m**). **n–q,** Representative immunoblots of UCP1 and β-actin in IBAT (**n**) and ingWAT (**p**), and quantification of total UCP1 content in one IBAT lobe (**o**) and one ingWAT pad (**q**). The mean value for IBAT of control mice was set to 1.0, and all values are expressed relative to this value. **r,** Total UCP1 per mouse, calculated as the sum of UCP1 content in IBAT and ingWAT. **s,** Correlation between CL-induced increase in oxygen consumption (from g) and total UCP1 content (from r). Each symbol represents one mouse. Values are means ± SEM. Asterisks indicate significant differences between control and UCP1^Myf5cKO^ mice. P* < 0.05, P** < 0.01, ***P < 0.001 (two-tailed unpaired Student’s t-test).

To validate the model, adipose tissues from mice acclimated to 4 °C for 1–2 months (Figure 1b) were analyzed by immunohistochemistry. This condition imposes high thermogenic demand and promotes the emergence of beige adipocytes. UCP1 immunostaining identified thermogenic adipocytes, whereas perilipin immunostaining visualized the entire adipocyte population. In UCP1^Myf5cKO^ mice, selective deletion of UCP1 in Myf5-derived adipocytes enabled UCP1-positive adipocytes to be identified as of non-Myf5 origin (Figure 1c,d). Interscapular brown adipose tissue (IBAT) and inguinal white adipose tissue (ingWAT) were selected as representative brown and beige adipose depots, respectively.

In IBAT of control mice, UCP1-positive adipocytes were distributed uniformly throughout the tissue (Figure 1c, upper panels). In contrast, cold-acclimated UCP1^Myf5cKO^ mice showed a marked phenotypic change, with the majority of adipocytes lacking UCP1 expression, although UCP1-positive adipocytes were not completely absent (Figure 1c, lower panels). These UCP1-positive adipocytes were detected sporadically, often in island-like clusters, suggestive of a clonal origin. Their non-Myf5 origin is consistent with previous reports that a small adipocyte population within IBAT is not of Myf5 origin^9^ and may correspond, at least in part, to the TRPV1-lineage of thermogenic adipocytes^19^. The conclusion that the remaining UCP1-positive adipocytes were not of Myf5 origin was supported in triple-mutant Myf5-Cre^+/-^; UCP1^fl/fl^; Rosa-Yfp^+/+^ mice, where YFP and UCP1 expression were mutually exclusive in adult tissue (Figure S2a-b).

UCP1-positive adipocytes were present in ingWAT of both control and UCP1^Myf5cKO^ mice (Figure 1d), consistent with the non-Myf5 origin of beige adipocytes. Notably, ingWAT from UCP1^Myf5cKO^ mice appeared to contain a greater number of strongly UCP1-positive adipocytes arranged in larger islands (Figure 1d, lower panels), suggesting a compensatory increase in this depot in response to the marked reduction of UCP1 expression in brown adipose tissue

Together, these phenotypes confirmed the selective deletion of UCP1 in adipocytes of myogenic origin while preserving UCP1 expression in adipocytes of non-myogenic origin, thereby validating the UCP1^Myf5cKO^ model.

### Cold-acclimated UCP1^Myf5cKO^ mice display an unaltered thermogenic response to CL316,243 despite markedly diminished UCP1 content

Because cold-acclimation-recruited adrenergically-induced nonshivering thermogenesis is UCP1-dependent^13,14,20^, the marked reduction of UCP1 expression in IBAT makes the UCP1^Myf5cKO^ model well suited for assessing whether beige adipocytes alone can support nonshivering thermogenesis during cold acclimation.

Nonshivering thermogenic capacity was assessed in conscious mice by measuring oxygen consumption following administration of the β3-adrenergic agonist CL316,243 (CL). Because this response reflects the heat-generating capacity of UCP1-containing adipocytes, it has been considered proportional to total UCP1 content^21^. CL rapidly increased oxygen consumption (Figure 1e), remarkably similarly in control and UCP1^Myf5cKO^ mice. Because of the long-lasting action of CL, values obtained 2–4 h after injection, when oxygen consumption had reached a stable plateau, were used to quantify the thermogenic response (Figure 1f). The CL-induced increase in oxygen consumption did not differ between control and UCP1^Myf5cKO^ mice (Figure 1g). Control experiments confirmed that the sustained increase in oxygen consumption reflected the pharmacological action of CL rather than transient injection-induced stress, and the increase in both genotypes was accompanied by a rapid decline in RER to ∼0.7, consistent with increased lipid utilization (Figure S2e; Figure S6e-g).

This unaltered thermogenic capacity in UCP1^Myf5cKO^ mice may suggest that the marked reduction of UCP1 in brown adipose tissue may be compensated by increased UCP1 content in beige adipose tissue. The increased abundance of UCP1-positive beige adipocytes observed by immunohistochemistry is consistent with this possibility. We therefore assessed basic biochemical characteristics and quantified UCP1 protein levels in the animals used for thermogenic measurements, focusing on IBAT and ingWAT.

UCP1 ablation in Myf5-originating brown adipocytes did not affect IBAT wet weight (Figure 1h) but markedly reduced total IBAT protein content (Figure 1i). In contrast, both wet weight and total protein content were increased in ingWAT from cold-acclimated UCP1^Myf5cKO^ mice (Figure 1k,l). Together with reduced protein density in IBAT and a tendency toward increased protein density in ingWAT (Figure 1j,m), these findings are consistent with brown-fat atrophy and enhanced beige-fat recruitment.

In agreement with the immunohistochemical observations, UCP1 protein levels in IBAT were markedly reduced in UCP1^Myf5cKO^ mice (Figure 1n and Figure S2f), being undetectable in some samples and substantially lower in others. Because total UCP1 content is considered the parameter most closely related to tissue thermogenic capacity^21,22^, we calculated total IBAT UCP1 content (Figure 1o) by multiplying UCP1 protein levels (Figure S2f) with total tissue protein content (Figure 1i), revealing a profound reduction in the amount of UCP1 available for thermogenesis.

In contrast, total UCP1 content in ingWAT was increased (Figure 1q), consistent with compensatory recruitment of this beige depot. However, even when this increase was taken into account, the combined UCP1 content of IBAT and ingWAT remained much lower than in controls, with only ∼30 % remaining (Figure 1r). Thus, the substantial reduction in total UCP1 content did not translate into impaired thermogenic capacity in cold-acclimated UCP1^Myf5cKO^ mice. Correlation analysis revealed no relationship between total UCP1 content per mouse (Figure 1r) and the CL-induced increase in oxygen consumption (Figure 1g) in these cold-acclimated mice (Figure 1s).

These findings challenged a simple quantitative relationship between UCP1 content and thermogenic capacity and prompted further experimental analyses to define the physiological basis of the observed phenotype.

### Whole-body UCP1 content remains markedly reduced in UCP1^Myf5cKO^ mice despite compensatory recruitment of adipose depots of non-Myf5 origin

Because thermogenic capacity in UCP1^Myf5cKO^ mice was maintained despite markedly reduced total UCP1 content in IBAT and ingWAT together, we examined additional adipose depots that could contribute substantially to whole-body UCP1 content^1^: perirenal BAT (pr BAT), of non-Myf5 origin; cervical BAT (cBAT), of mixed origin; and axillary BAT (aBAT), of predominantly Myf5 origin^9^. Mice were acclimated to thermoneutrality or cold for ∼2 months (Figure 2a).

**Figure 2.**
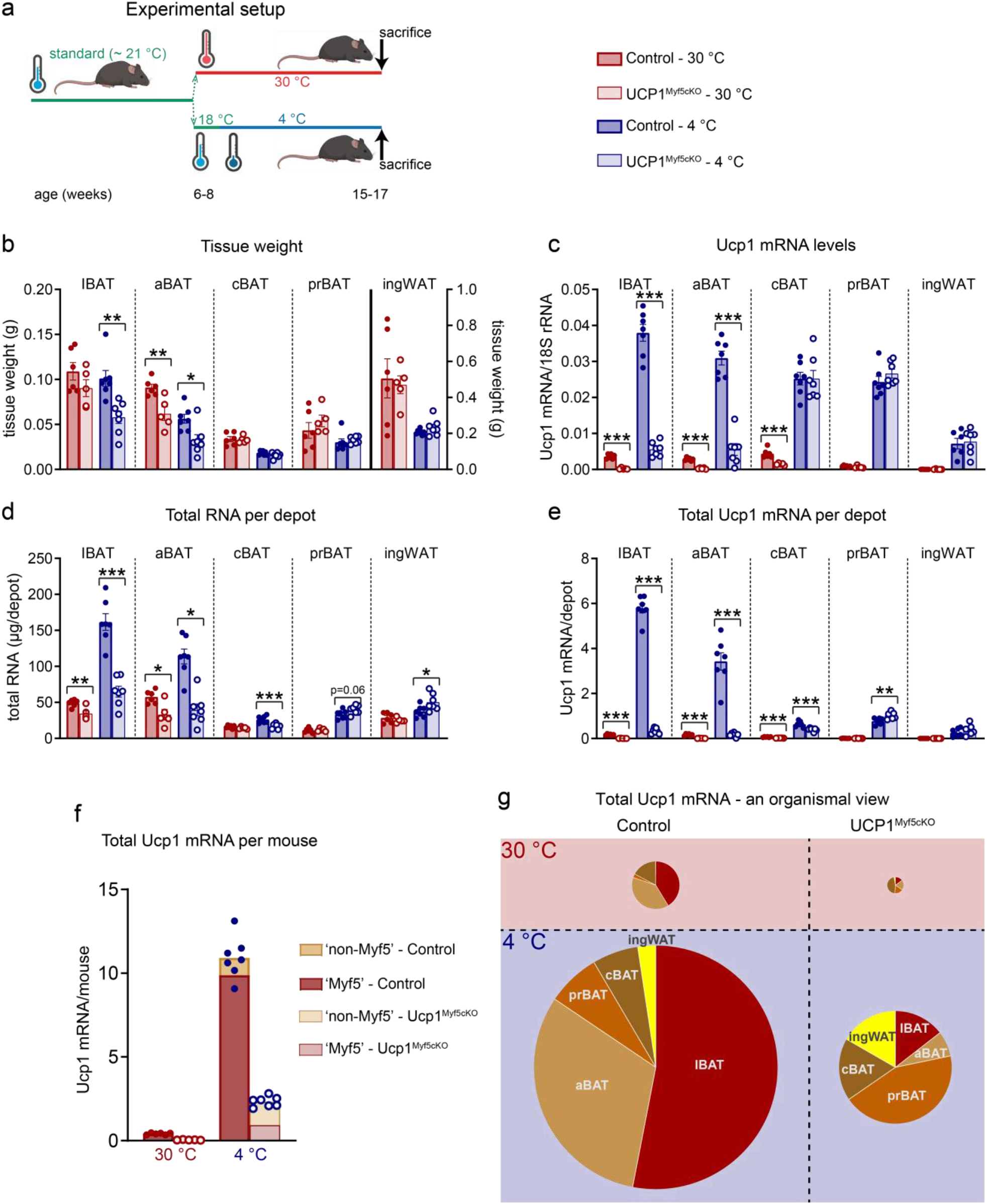
In cold-acclimated UCP1^Myf5cKO^ mice, total Ucp1 mRNA content is markedly reduced despite compensatory recruitment of non-Myf5-derived adipose depots. **a,** Experimental design. Control and UCP1^Myf5cKO^ mice were acclimated to 30 °C (control, n = 6; UCP1^Myf5cKO^, n = 5) or 4 °C (control, n = 7; UCP1^Myf5cKO^, n = 7). The following adipose depots were analysed: interscapular brown adipose tissue (IBAT), axillary BAT (aBAT), cervical BAT (cBAT), perirenal BAT (prBAT), and inguinal white adipose tissue (ingWAT). **b,** Wet weight of the indicated adipose depots. **c,** Ucp1 mRNA levels normalized to 18S rRNA (Figure S3a). **d,** Total RNA content per depot. **e,** Total Ucp1 mRNA content per depot, calculated by multiplying Ucp1 mRNA levels **(c)** by total RNA content **(d)**. **f,** Total Ucp1 mRNA content per mouse, calculated as the sum of all examined adipose depots. Bar segments indicate contributions from Myf5-derived and non-Myf5-derived depots. **g,** Relative contribution of each adipose depot to total Ucp1 mRNA content in mice acclimated to thermoneutrality or cold. Each symbol represents one mouse. Values are means ± SEM. Asterisks indicate significant differences between control and UCP1^Myf5cKO^ mice. *P < 0.05, **P < 0.01, ***P < 0.001 (two-tailed unpaired Student’s t-test).

Cold acclimation generally decreased the wet weight of all examined adipose depots (Figure 2b). Genotype effects were confined to predominantly Myf5-derived depots, with reduced IBAT and aBAT weight in UCP1^Myf5cKO^ mice.

Because of the large number of samples (five tissues across two genotypes and two temperature conditions), UCP1 expression was quantified at the mRNA level that accurately reflects total UCP1 protein under steady-state conditions^2^. Cold acclimation robustly increased UCP1 mRNA expression in all depots (Figure 2c), with particularly large relative increases in prBAT and ingWAT, where basal expression is low^1,23^. At both acclimation temperatures, UCP1 mRNA expression was reduced in Myf5-derived depots (IBAT and aBAT) of UCP1^Myf5cKO^ mice, while remaining largely unchanged in non-Myf5 depots (prBAT and ingWAT); cBAT showed a reduction only under thermoneutral conditions.

To estimate the contribution of each depot to thermogenic capacity, UCP1 mRNA content per depot was calculated by multiplying UCP1 mRNA levels (Figure 2c) by total RNA per depot (Figure 2d). Total RNA content increased with cold acclimation in all examined depots. Analysis of UCP1 mRNA content at the level of individual depots (Figure 2e) showed that IBAT and aBAT were the major contributors to total UCP1 content in control mice. In UCP1^Myf5cKO^ mice, UCP1 mRNA content was reduced in Myf5-derived depots (IBAT and aBAT) but increased in non-Myf5-derived depots (prBAT and ingWAT).

To integrate these depot-specific observations, total UCP1 mRNA per mouse was calculated and presented as absolute values (bar graphs), with fractions of Myf5 and non-Myf5 origin indicated (Figure 2f). The relative contribution of each depot is shown in Figure 2g.

As evident from this organismal-level analysis, a compensatory increase in UCP1 content was observed in adipose depots of non-Myf5 origin in cold-acclimated UCP1^Myf5cKO^ mice. Nevertheless, total UCP1 content in cold-acclimated UCP1^Myf5cKO^ mice remained only ∼20 % of control levels (Figure 2f,g), despite a greater relative cold-induced recruitment (∼50-fold) than in control mice (∼25-fold). Thus, compensatory recruitment of adipose depots of non-Myf5 origin was insufficient to restore whole-body UCP1 content.

### No evidence for compensatory recruitment of UCP1-independent thermogenic pathways

Because compensatory recruitment of non-Myf5 adipose depots could not account for the preserved thermogenic capacity of UCP1^Myf5cKO^ mice, we next examined whether suggested UCP1-independent thermogenic pathways were compensatorily recruited. We measured the expression of key components of futile creatine cycling^24–26^ (Figure S3b), futile calcium cycling^27^ (Figure S3c), futile lipid cycling^28–30^ (Figure S3d), ADP/ATP carrier-mediated uncoupling^31^ (Figure S3e), and N-acyl amino acid-mediated uncoupling^32^ (Figure S3f). Gene expression was analyzed in IBAT and ingWAT, representing predominantly Myf5- and non-Myf5-derived adipose depots, respectively.

In IBAT from cold-acclimated UCP1^Myf5cKO^ mice, mRNA expression of several genes associated with proposed UCP1-independent thermogenic pathways, including Ckb, Gk, and Dgat1, was increased. However, owing to pronounced tissue atrophy, total transcript content was generally reduced when assessed at the whole-depot level (Figure S3, bottom panels).

In ingWAT, no genotype differences were detected in the expression levels or total transcript content of genes associated with these pathways. Thus, no evidence was obtained for compensatory recruitment of proposed UCP1-independent thermogenic mechanisms in the beige depot.

Together, these observations provide little support for a physiologically relevant contribution of proposed UCP1-independent thermogenic pathways to the preserved thermogenic capacity of cold-acclimated UCP1^Myf5cKO^ mice.

### Cold-acclimation–recruited adrenergically induced nonshivering thermogenesis is largely absent in global UCP1-KO mice but preserved in UCP1^Myf5cKO^ mice

Because no evidence for compensatory recruitment of UCP1-independent thermogenic pathways was found, we next asked whether the amount of UCP1 present was sufficient to sustain adaptive nonshivering thermogenesis in UCP1^Myf5cKO^ mice. To test this, thermogenic capacity was directly compared between the UCP1^Myf5cKO^ mice, that express low levels of UCP1, and global UCP1-KO mice, that completely lack UCP1.

Global UCP1-KO mice sustain high oxygen consumption under cold conditions^13,33,34^, yet recruit only a very limited adrenergically induced thermogenic response during cold acclimation^13,35^. Thus, in these mice adaptive nonshivering thermogenesis is largely absent, with shivering constituting the major source of heat production^13^. Global UCP1-KO mice therefore provide a framework to determine whether the preserved thermogenic response of UCP1^Myf5cKO^ mice is mediated by the low levels of UCP1 present.

To maximize the contrast between nonrecruited and fully recruited thermogenic capacity, UCP1^Myf5cKO^, global UCP1-KO, and corresponding control mice were acclimated to thermoneutrality (30 °C) or cold (4 °C) for ∼2 months. Diurnal energy expenditure was measured by indirect calorimetry at the corresponding temperatures. All groups displayed normal diurnal variation, with higher energy expenditure during the active phase (Figure 3a,b). Energy expenditure was largely unaffected by complete or partial UCP1 ablation and was approximately threefold higher in the cold-acclimated mice remaining in the cold (Figure 3c,d). A minor reduction was observed in cold-acclimated global UCP1-KO mice, consistent with their lower lean mass (Figure S4a).

**Figure 3.**
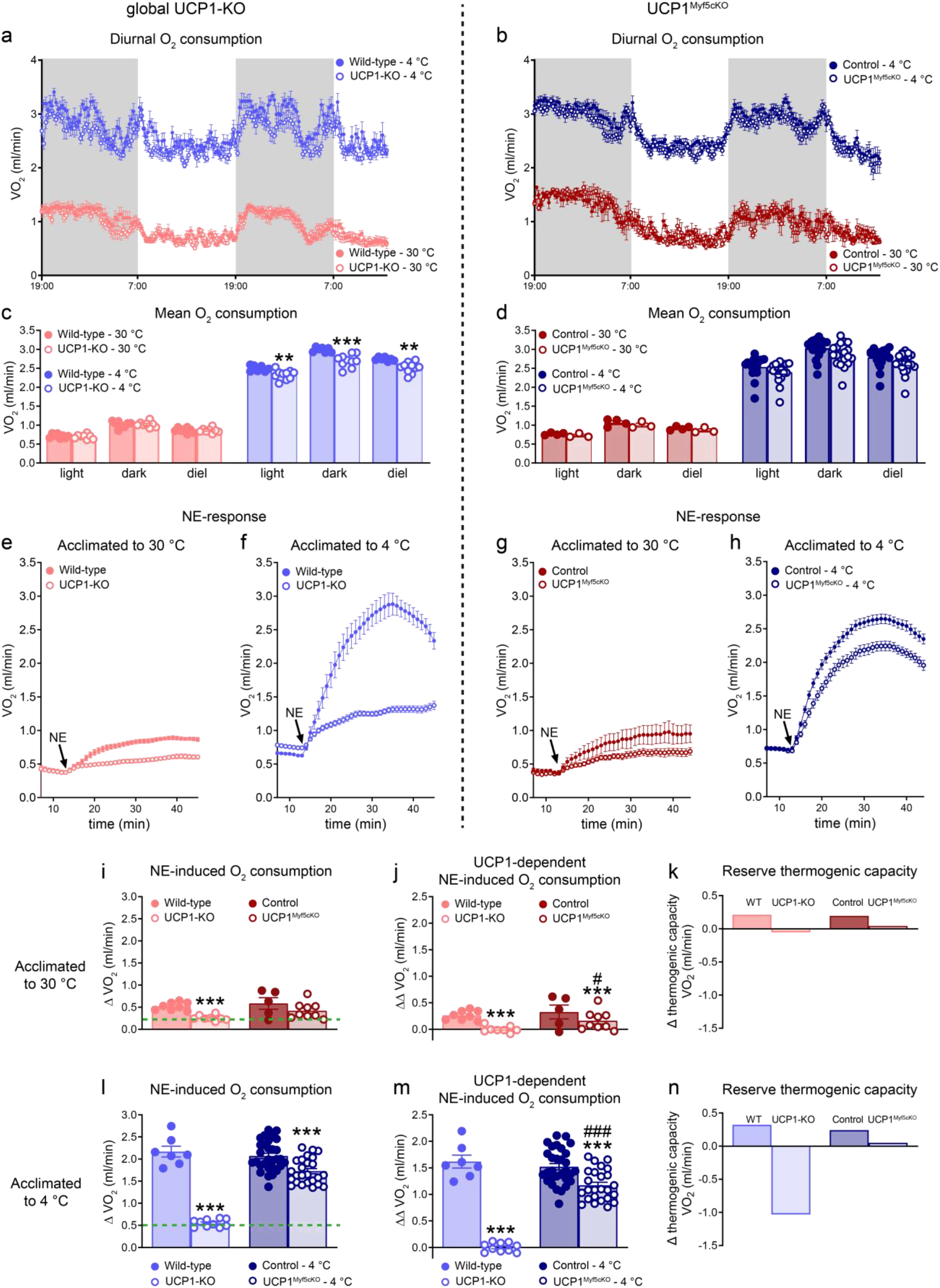
UCP1 is essential but not rate-limiting for cold acclimation-recruited adrenergically induced nonshivering thermogenesis. Global UCP1-KO mice and UCP1^Myf5cKO^ mice, together with their respective controls, were acclimated to 30 °C or 4 °C as shown in Figure 2a. **a,b,** Diurnal oxygen consumption measured by indirect calorimetry at the corresponding acclimation temperature at the end of the acclimation period. Gray shading indicates the dark phase. **a,** Wild-type (30 °C, n = 8; 4 °C, n = 8) and UCP1-KO mice (30 °C, n = 8; 4 °C, n = 10). **b,** Control (30 °C, n = 4; 4 °C, n = 23) and UCP1^Myf5cKO^ mice (30 °C, n = 3; 4 °C, n = 19). **c,d,** Mean daytime, nighttime, and 24-h (diel) oxygen consumption calculated from a and b. **e–h,** Thermogenic capacity assessed by the norepinephrine (NE) test in pentobarbital-anesthetized mice. **e,** Wild-type (n = 8) and UCP1-KO mice (n = 8) acclimated to 30 °C. **f,** Wild-type (n = 7) and UCP1-KO mice (n = 10) acclimated to 4 °C. **g,** Control (n = 5) and UCP1^Myf5cKO^ mice (n = 9) acclimated to 30 °C. **h,** Control (n = 30) and UCP1^Myf5cKO^ mice (n = 25) acclimated to 4 °C. **i,l,** NE-induced increase in oxygen consumption, calculated for each mouse as maximal oxygen consumption after NE injection minus baseline oxygen consumption before injection. **i,** Mice acclimated to 30 °C. **l,** Mice acclimated to 4 °C. **j,m,** UCP1-dependent component of the NE response, calculated by subtracting the mean NE-induced oxygen consumption of UCP1-KO mice at the corresponding acclimation temperature (green dashed line in **i** and **l**) from individual value for each mouse (including UCP1-KO mice; thus, for UCP1-KO mice, this yields 0). **j,** Mice acclimated to 30 °C. **m,** Mice acclimated to 4 °C. **k,n,** Estimated reserve nonshivering thermogenic capacity at 30 °C (**k**) and 4 °C (**n**), calculated as the difference between average maximal NE-induced oxygen consumption (Figure 3e-h and Figure S3c,d) and average energy expenditure during the light phase (Figure 3c,d). Positive values indicate reserve nonshivering thermogenic capacity, whereas negative values indicate that maximal nonshivering thermogenesis is insufficient to meet the estimated thermogenic demand. Each symbol represents one mouse. Values are means ± SEM. Asterisks indicate differences between each genetically modified strain and its corresponding control (*P < 0.05, **P < 0.01, ***P < 0.001); hashtags indicate differences between UCP1-KO and UCP1^Myf5cKO^ mice (^#^P < 0.05, ^###^P < 0.001) (two-tailed unpaired Student’s t-test).

To assess nonshivering thermogenic capacity, oxygen consumption was measured following norepinephrine (NE) administration. Unlike the β3-adrenergic agonist CL, NE activates multiple adrenergic receptor subtypes and therefore provides a more physiological estimate of adrenergically induced thermogenesis. Measurements were performed under anesthesia to eliminate the confounding thermogenic effects of injection-induced stress^14^.

In thermoneutral wild-type mice, NE increased metabolic rate by ∼130 %, whereas global UCP1-KO mice showed a smaller response of ∼70 % (Figure 3e), representing the UCP1-independent component of the NE response.

Cold acclimation markedly enhanced the NE response in wild-type mice (>4-fold above baseline; Figure 3f); in global UCP1-KO mice, the response again reached only ∼70 % above baseline. Cold acclimation-recruited nonshivering thermogenesis was thus largely UCP1 dependent, as expected^13,14,35^.

Under thermoneutral conditions, UCP1^Myf5cKO^ mice displayed only a modest reduction in the NE response compared with controls (Figure 3g,i). After subtraction of the average response observed in global UCP1-KO mice (indicated by the green dashed line in Figure 3i), representing the UCP1-independent component, a small but detectable UCP1-dependent thermogenic response remained (Figure 3j).

After cold acclimation, UCP1^Myf5cKO^ mice displayed a robust NE response (∼250 % above baseline), only modestly lower than that of control mice (∼300 %) and far greater than that of global UCP1-KO mice (∼70 %) (Figure 3h,l). Correspondingly, the calculated UCP1-dependent component of thermogenesis remained largely preserved (Figure 3m). Thus, mice retaining only ∼20 % of total UCP1 content exhibited ∼80 % of the thermogenic capacity observed in control mice.

To estimate whether the nonshivering thermogenic capacity recruited during cold acclimation was sufficient to defend body temperature in the cold, we calculated the difference between average maximal NE-induced oxygen consumption (Figure 3f,h and Figure S4d) and average energy expenditure during the light phase in the cold (Figure 3c,d). The latter was used as an approximation of the minimal thermogenic demand required to maintain body temperature at 4 °C. Positive values indicate reserve thermogenic capacity, whereas negative values indicate insufficient nonshivering thermogenic capacity.

Based on Figure 3n, global UCP1-KO mice displayed a pronounced nonshivering thermogenic deficit, consistent with a requirement for additional heat production, most likely through shivering. Corresponding control mice retained a clear reserve thermogenic capacity. Importantly, UCP1^Myf5cKO^ mice possessed just sufficient nonshivering thermogenic capacity to maintain body temperature in the cold. Thus, these mice would not be expected to require engagement of shivering thermogenesis. At thermoneutrality, where no additional thermogenesis is required for maintenance of body temperature, both global UCP1-KO mice and UCP1^Myf5cKO^ mice were approximately in energetic balance, whereas corresponding control mice retained a small reserve thermogenic capacity (Figure 3k).

Together, these findings demonstrate that cold acclimation-recruited adaptive nonshivering thermogenesis requires UCP1 but is not constrained by its abundance. Maximal thermogenic capacity is achieved with only a fraction of the UCP1 normally present.

### Loss of one functional Ucp1 allele neither elicits compensatory adaptations within IBAT nor compromises cold acclimation-recruited nonshivering thermogenesis

Previous work using mice heterozygous for a global Ucp1 deletion demonstrated that UCP1 protein levels in IBAT are approximately 50 % of wild-type levels at room temperature^36^, indicating that loss of one *Ucp1* allele does not elicit compensatory upregulation of expression from the remaining *Ucp1* allele. Because this global knockout model was generated from the same EUCOMM *Ucp*^1tm1a^ allele and involves deletion of the same *Ucp1* gene region as in the present study, this observation is directly relevant to the present conditional model. We therefore examined cold-acclimated mice heterozygous for the conditional Ucp1 deletion (Myf5-Cre^+/-^; Ucp1^fl/+^; hereafter referred to as Het-UCP1^Myf5cKO^ mice) to determine whether loss of one functional *Ucp1* allele elicits compensatory adaptations under *high* thermogenic demand. If compensatory adaptations were elicited, Ucp1 transcript levels would be expected to exceed the ∼50 % level expected from gene dosage alone and/or sympathetic innervation of IBAT would be expected to increase. Conversely, the absence of these adaptations would suggest that the remaining UCP1 content is sufficient under these conditions and therefore not functionally limiting.

As shown in Figure 4a, UCP1 mRNA content in IBAT of cold-acclimated Het-UCP1^Myf5cKO^ mice remained at approximately 50 % of control levels. Thus, even under conditions of markedly increased thermogenic demand, expression from the remaining *Ucp1* allele was not compensatorily upregulated. Because both *Ucp1* alleles remain intact in adipose depots not derived from Myf5-positive progenitors, compensatory increases in UCP1 expression could, in principle, also occur in these depots. However, total UCP1 mRNA content per mouse was still ∼40 % lower than in controls (Figure 4b), indicating that such compensation did not occur.

**Figure 4.**
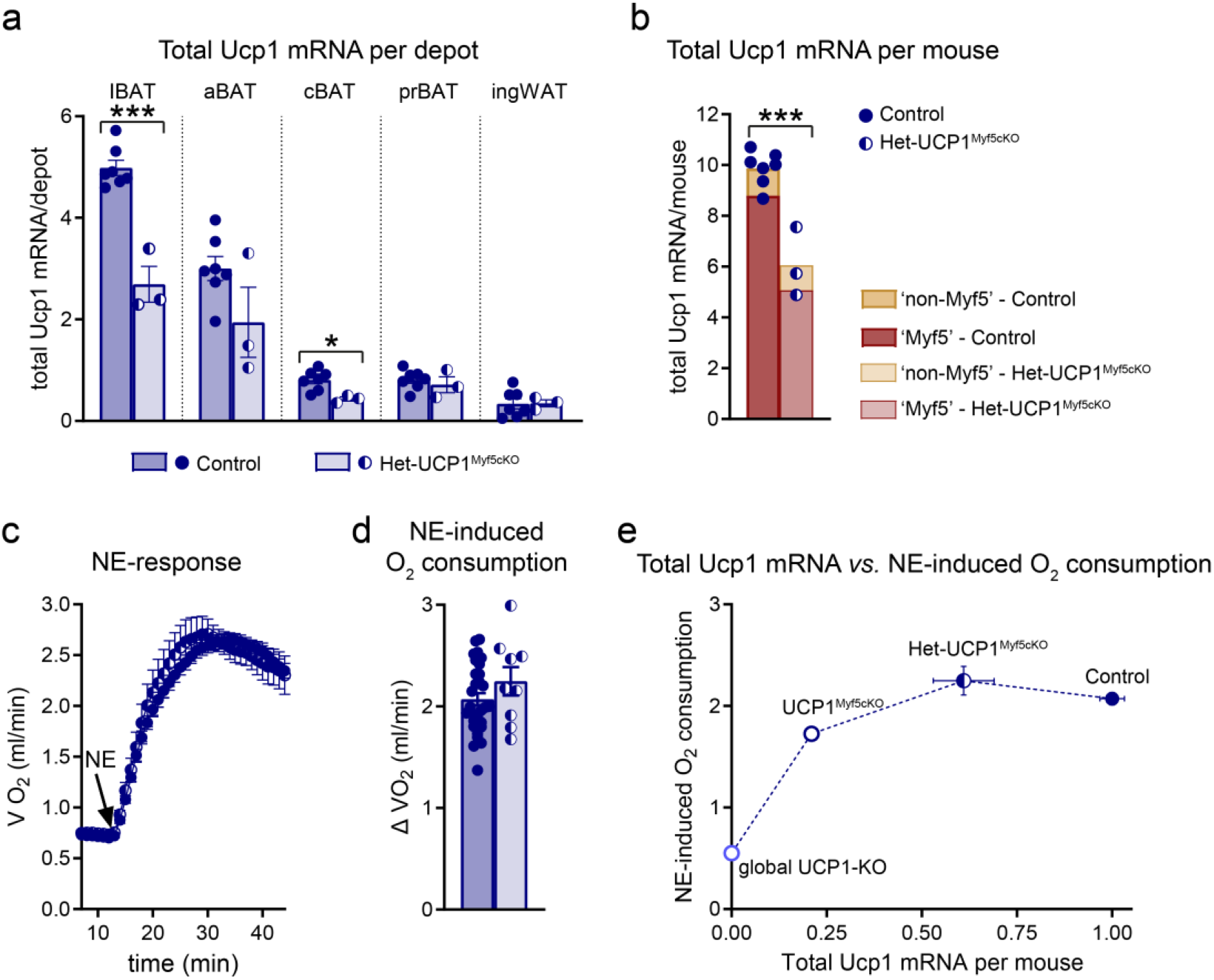
Cold acclimation-recruited nonshivering thermogenesis is fully preserved despite an approximately 50 % reduction in UCP1 content. **a,b,** Control (n = 7) and heterozygous Het-UCP1^Myf5cKO^ (n = 3) mice were acclimated to 4 °C. Control mice are the same as those shown in Figure 2 and are reanalyzed in parallel with Het-UCP1^Myf5cKO^ mice. The same adipose depots as in Figure 2 were analyzed. **a,** Total Ucp1 mRNA content per depot, calculated by multiplying Ucp1 mRNA levels (Figure S5a) by total RNA content (Figure S5b). **b,** Total Ucp1 mRNA content per mouse, calculated as the sum of all examined adipose depots. Bar segments indicate contributions from Myf5-derived and non-Myf5-derived depots. **c,d,** Control (n = 30) and Het-UCP1^Myf5cKO^ mice (n = 9) were acclimated to 4 °C. Control mice are the same as those shown in Figure 3h,j and are replotted for comparison with Het-UCP1^Myf5cKO^ mice. **c,** Oxygen consumption before and after NE injection. **d**, NE-induced increase in oxygen consumption, calculated as maximal oxygen consumption after NE injection minus baseline oxygen consumption before NE injection. **e,** Relationship between whole-body Ucp1 mRNA content and NE-induced thermogenic response. Group mean values are shown because the two variables were measured in independent cohorts. The line is shown to guide the eye. Each symbol represents one mouse. Values are means ± SEM. Asterisks indicate significant differences between control and Het-UCP1^Myf5cKO^ mice. P* < 0.05, ***P < 0.001 (two-tailed unpaired Student’s t-test).

Despite this reduction in total UCP1 content, thermogenic capacity remained unchanged in cold-acclimated Het-UCP1^Myf5cKO^ mice (Figure 4c,d). These mice also retained reserve thermogenic capacity comparable to that of controls (not shown). Thus, this substantial reduction in total UCP1 content did not compromise cold acclimation-recruited thermogenic capacity, indicating that UCP1 is present in excess of the amount required to sustain maximal thermogenic output in cold-acclimated mice.

To further illustrate the relationship between total UCP1 content and thermogenic capacity, mean norepinephrine-induced thermogenic responses across genotypes (Figures 3l and 4d) were plotted against mean total UCP1 mRNA content (Figures 2f and 4b). Despite marked differences in UCP1 mRNA content, control, UCP1^Myf5cKO^, and Het-UCP1^Myf5cKO^ mice clustered closely together with similarly high thermogenic responses, whereas global UCP1-KO mice were clearly separated by a markedly reduced response (Figure 4e). Although this comparison is based on group means obtained from independent cohorts rather than individual correlations, it provides independent support for the conclusion that UCP1 is essential but not rate-limiting for adaptive nonshivering thermogenesis in cold-acclimated mice.

We recently showed that complete UCP1 deficiency is associated with markedly increased sympathetic innervation of IBAT, reflecting a feedback response to the impaired heat production^37^, and we here examined whether a similar response would exist in Het-UCP1^Myf5cKO^ mice. Sympathetic innervation was assessed by tyrosine hydroxylase (TH) immunostaining (Figure 5a–l). TH immunoreactivity was similar in control and Het-UCP1^Myf5cKO^ IBAT (Figure 5b,f) but markedly increased in IBAT of UCP1^Myf5cKO^ mice (Figure 5j), indicating enhanced sympathetic innervation only following profound UCP1 depletion. As expected, UCP1 immunostaining in UCP1^Myf5cKO^ IBAT exhibited a mosaic pattern, with most adipocytes lacking UCP1 and only a minority remaining positive (Figure 5i,l). In contrast, UCP1 staining was uniformly distributed in control and heterozygous (Het-UCP1^Myf5cKO^) IBAT but appeared less intense in heterozygous tissue (Figure 5a,e).

**Figure 5.**
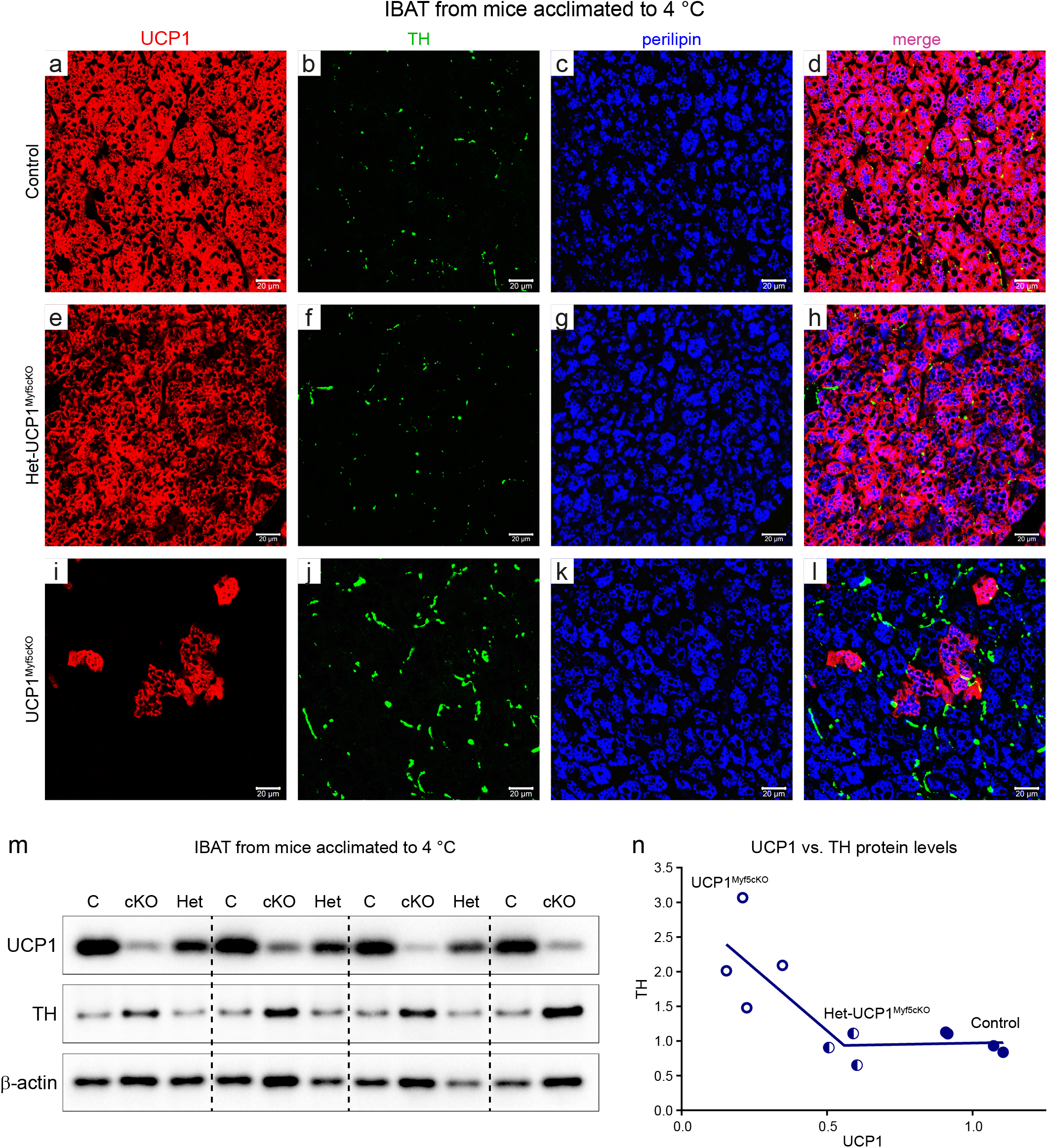
Approximately half of the normal UCP1 content does not induce compensatory sympathetic remodeling in IBAT. **a–l**, Representative confocal images of IBAT from control, Het-UCP1^Myf5cKO^, and UCP1^Myf5cKO^ mice acclimated to 4 °C. Sections were stained for UCP1 (red; **a,e,i**), tyrosine hydroxylase (TH; green; **b,f,j**), and perilipin (blue; **c,g,k**). **d,h,l,** Merged images. Scale bar, 20 μm. **m,** Representative immunoblots of UCP1, TH, and β-actin in IBAT from control, Het-UCP1^Myf5cKO^, and UCP1^Myf5cKO^ mice acclimated to 4 °C. **n,** Relationship between UCP1 and TH protein levels based on immunoblot quantification in m. Mean UCP1 and TH protein levels in control IBAT (n = 4) were set to 1.0, and individual values for control, Het-UCP1^Myf5cKO^ (n = 3), and UCP1^Myf5cKO^ (n = 4) mice are expressed relative to these mean control values. Each symbol represents one mouse. Values are means ± SEM. Data were fitted using segmental linear regression.

These observations were confirmed by immunoblotting (Figure 5m). UCP1 protein levels were reduced to approximately 50 % of control levels in IBAT of Het-UCP1^Myf5cKO^ mice and were markedly reduced in IBAT of UCP1^Myf5cKO^ mice. In contrast, TH protein levels were increased only in UCP1^Myf5cKO^ mice, whereas control and Het-UCP1^Myf5cKO^ mice displayed similar levels. This relationship was further illustrated by plotting TH against UCP1 protein levels for individual IBAT samples (Figure 5n).

Together, these molecular, structural, and functional analyses (Figures 4 and 5) demonstrate that a ∼50 % reduction in UCP1 abundance neither elicits compensatory adaptations within IBAT nor compromises cold acclimation-recruited thermogenic capacity. These findings indicate that the remaining UCP1 is sufficient to sustain normal thermogenic function within IBAT. Enhanced sympathetic innervation was observed only after profound UCP1 depletion in UCP1^Myf5cKO^ mice, indicating that compensatory sympathetic remodeling is triggered only by substantial loss of UCP1.

### A one-third reduction in mitochondrial UCP1 does not reduce UCP1-dependent respiration in fully recruited brown-fat mitochondria: evidence for a reserve UCP1 capacity

Our earlier studies of brown-fat mitochondria isolated from mice acclimated to different temperatures showed that UCP1-dependent respiration is proportional to UCP1 abundance at lower levels of thermogenic recruitment but approaches maximal respiratory capacity in fully recruited brown-fat mitochondria from cold-acclimated mice^3,6,38^. These observations can be interpreted as suggesting that fully recruited mitochondria may contain more UCP1 than is required to support maximal UCP1-dependent respiration. Direct testing of this hypothesis required brown-fat mitochondria with a similar degree of thermogenic recruitment but reduced UCP1 abundance. Cold-acclimated Het-UCP1^Myf5cKO^ mice provided such a model. We therefore examined respiratory function in brown-fat mitochondria isolated from cold-acclimated control and Het-UCP1^Myf5cKO^ mice.

As shown in Figure 6a, substrate addition (pyruvate, malate and octanoyl-L-carnitine) resulted, as expected, in a spontaneous high rate of oxygen consumption (thermogenesis) that was inhibited by GDP. The GDP-inhibitable component of respiration is mediated by UCP1^38–42^. ADP induced only a small increase in respiration, reflecting low oxidative phosphorylation capacity. FCCP was used to reveal maximal respiratory capacity. Mitochondria from Het-UCP1^Myf5cKO^ mice displayed an almost identical respiratory profile to those from control mice (Figure 6a), and quantitative analysis of multiple independent mitochondrial preparations showed no overall effect of genotype (Figure 6b).

**Figure 6.**
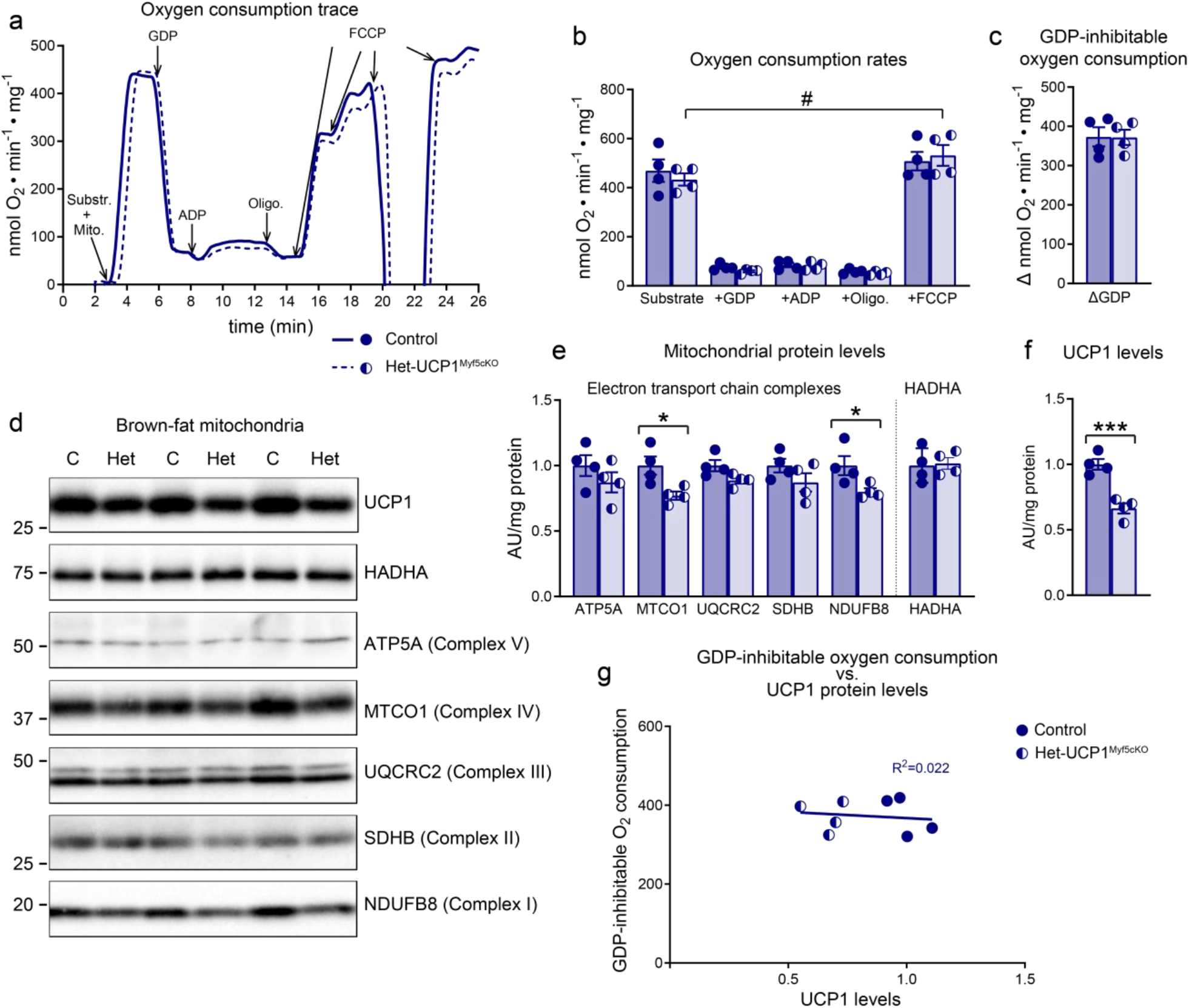
A one-third reduction in mitochondrial UCP1 does not reduce UCP1-dependent respiration in brown-fat mitochondria from cold-acclimated mice. Brown-fat mitochondria were isolated from control and heterozygous Het-UCP1^Myf5cKO^ mice acclimated to 4 °C. **a,** Representative traces of oxygen consumption in isolated brown-fat mitochondria. Additions were 0.125 mg mitochondria, substrate (5 mM malate + 5 mM pyruvate + 0.5 mM Octanoyl-L-carnitine), GDP (2 mM), ADP (450 μM), oligomycin (6 μg/ml), and FCCP (0.6 μM followed by up to three additions of 0.2 μM). **b,** Oxygen consumption rates compiled from experiments as shown in a (control, n = 4; Het-UCP1^Myf5cKO^, n = 4 independent mitochondrial preparations). **c,** The GDP-inhibitable component of oxygen consumption**. d–f,** Mitochondrial protein levels in the preparations analyzed in a–c. **d**, Representative immunoblots. **e,** Levels of representative respiratory-chain proteins and HADHA. **f,** UCP1 protein levels. Mean values in control mitochondria were set to 1.0, and values in all samples are expressed relative to these. **g,** GDP-inhibitable respiration (c) plotted against UCP1 abundance (f) for individual mitochondrial preparations. Each symbol represents one independent mitochondrial preparation. Values are means ± SEM. Asterisks indicate significant differences between control and Het-UCP1^Myf5cKO^ mice. *P < 0.05; ***P < 0.001 (two-tailed unpaired Student’s t-test). ^#^P < 0.05, FCCP-induced maximal respiration versus substrate-supported respiration (two-tailed paired Student’s t-test).

Immunoblot analysis of the same mitochondrial preparations showed a significant (∼33 %) reduction in mitochondrial UCP1 abundance in Het-UCP1^Myf5cKO^ mice. The abundance of respiratory-chain complexes I and IV was also modestly reduced, whereas the remaining respiratory-chain proteins and HADHA were comparable between genotypes (Figure 6d–f). Despite these modest reductions, FCCP-stimulated respiration did not differ between genotypes (Figure 6b). Importantly, under the experimental conditions used, an approximately one-third reduction in mitochondrial UCP1 abundance was not accompanied by a detectable reduction in UCP1-dependent respiration (Figure 6c). Thus, within the range of UCP1 abundance examined, reduced mitochondrial UCP1 abundance was compatible with preserved GDP-inhibitable respiration (Figure 6g).

Together, these results indicate that a substantial reduction in mitochondrial UCP1 abundance is compatible with preserved UCP1-dependent respiration in mitochondria isolated from brown adipose tissue of cold-acclimated mice. The mitochondrial findings complement the *in vivo* findings and support the interpretation that fully recruited brown-fat mitochondria contain more UCP1 than is required to sustain maximal UCP1-dependent respiration under the experimental conditions used. Because these experiments were performed in isolated mitochondria, this conclusion is restricted to intrinsic mitochondrial respiratory properties and cannot fully account for the regulation of thermogenesis *in vivo*.

### CL316,243-induced hypoxia identifies oxygen availability as a limiting factor for thermogenesis

The preceding experiments demonstrated that maximal thermogenic capacity is largely preserved despite a marked reduction in UCP1 abundance, indicating that UCP1 abundance is not the primary determinant of maximal thermogenic output. Because thermogenesis requires a continuous supply of oxygen to sustain mitochondrial respiration, previous observations that cold acclimation enhances vascularization and blood flow^43,44^, together with the finding that inhibition of angiogenesis impairs thermogenic capacity^45^, raised the possibility that oxygen (and nutrient) availability constrains maximal thermogenic capacity *in vivo*.

To address this, we developed an *in situ* hypoxia-labeling approach using pimonidazole. Cold-acclimated mice were transferred to thermoneutrality to suppress ongoing thermogenesis and subsequently stimulated with the long-acting β3-adrenergic agonist CL316,243 (CL), thereby maintaining thermogenic activity throughout the labeling period. The specificity of the protocol was verified by the absence of detectable hypoxia labeling in mice receiving pimonidazole alone (Figure S6a–d).

Importantly, this experimental design enabled simultaneous assessment of thermogenic capacity and tissue hypoxia in the same mice (Figure 7a–d). As in the norepinephrine experiments (Figure 3), cold-acclimated wild-type, UCP1^Myf5cKO^, and corresponding control mice exhibited robust CL-induced increases in oxygen consumption, whereas global UCP1-KO mice displayed only a limited response (Figure 7b,d). Injection of pimonidazole alone did not affect metabolic rate (Figure S6e,f), and CL administration rapidly lowered RER to ∼0.7, consistent with increased lipid utilization (Figure S6g).

**Figure 7.**
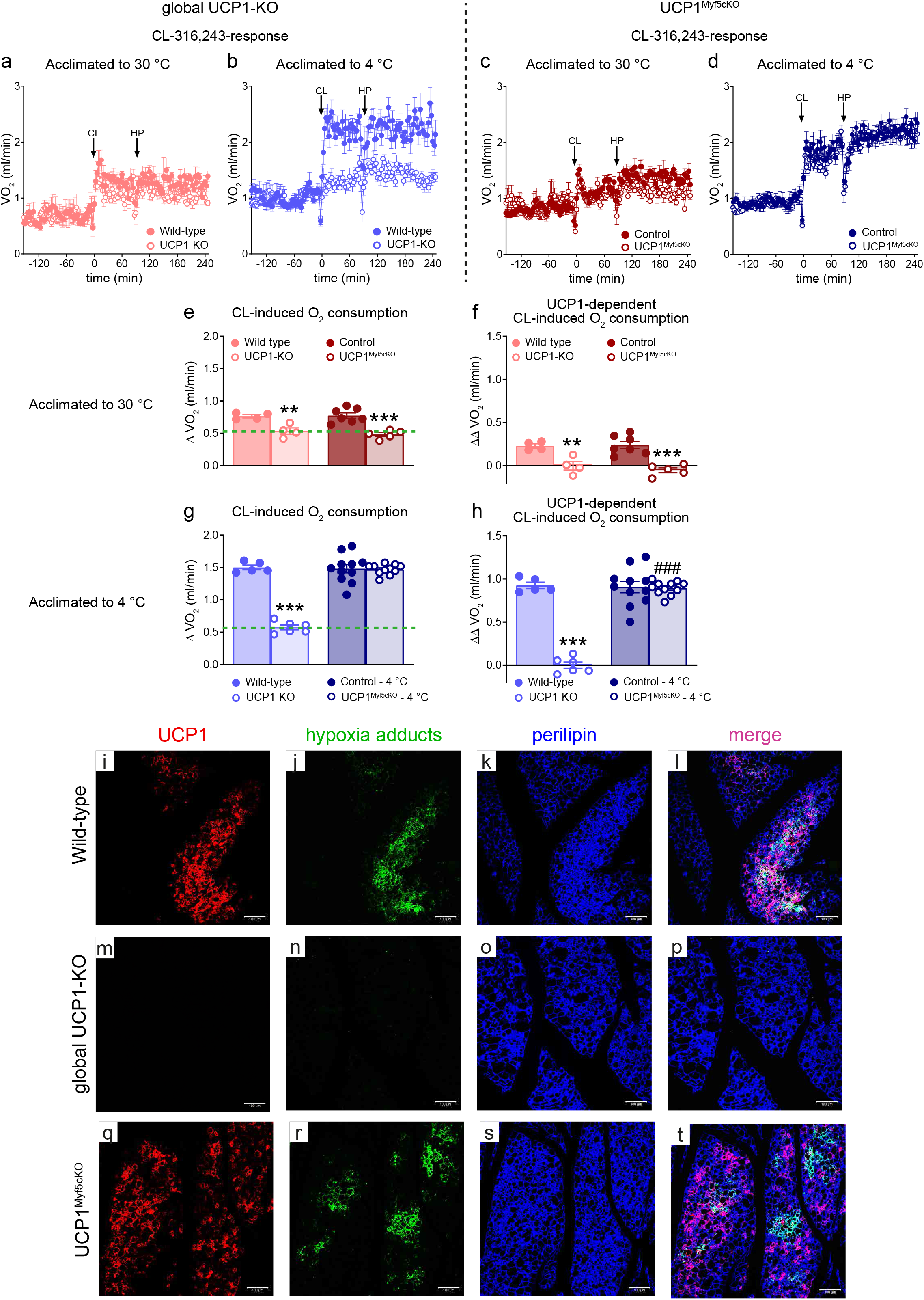
CL316,243-induced hypoxia identifies oxygen availability as a limiting factor for adaptive nonshivering thermogenesis in the cold. Global UCP1-KO mice and UCP1^Myf5cK^O mice, together with their respective controls, were acclimated to 30 °C or 4 °C as shown in Figure 2a. **a–d,** Thermogenic capacity assessed by indirect calorimetry following CL316,243 (CL) administration. Mice were housed overnight at 30 °C in the Promethion indirect calorimetry system, injected with CL (1 mg/kg), followed 90 min later by pimonidazole, and monitored for a further 2 h 30 min. **a,** Wild-type (n = 4) and UCP1-KO (n = 4) mice acclimated to 30 °C. **b,** Wild-type (n = 5) and UCP1-KO (n = 6) mice acclimated to 4 °C. **c,** Control (n = 7) and UCP1^Myf5cKO^ (n = 5) mice acclimated to 30 °C. **d,** Control (n = 11) and UCP1^Myf5cKO^ (n = 11) mice acclimated to 4 °C. **e,g,** CL-induced increase in oxygen consumption, calculated for each mouse as mean oxygen consumption during the final 2 h after CL administration minus baseline oxygen consumption before injection. **e,** Mice acclimated to 30 °C. **g,** Mice acclimated to 4 °C. **f,h,** UCP1-dependent component of the CL response, calculated by subtracting the mean CL-induced oxygen consumption of UCP1-KO mice at the corresponding acclimation temperature (green dashed line in **e** and **g**) from individual value for each mouse (including UCP1-KO mice; thus, for UCP1-KO mice, this yields 0). **f,** Mice acclimated to 30 °C. **h,** Mice acclimated to 4 °C. Each symbol represents one mouse. Values are means ± SEM. Asterisks indicate differences between each genetically modified strain and its corresponding control (**P < 0.01, ***P < 0.001); hashtags indicate differences between UCP1-KO and UCP1^Myf5cKO^ mice (^###^P < 0.001) (two-tailed unpaired Student’s t-test). **i–t,** Representative confocal images of ingWAT from cold-acclimated wild-type, UCP1-KO, and UCP1^Myf5cKO^ mice following CL and pimonidazole administration. Sections were stained for UCP1 (red; **i,m,q**), hypoxia adducts (green; **j,n,r**), and perilipin (blue**; k,o,s**). **l,p,t,** Merged images. Scale bar, 100 μm.

To estimate the UCP1-dependent component of thermogenesis, the average CL-induced oxygen consumption observed in global UCP1-KO mice at each acclimation temperature (Figure 7e,g, green dashed lines) was subtracted from the corresponding individual values. This analysis revealed a marked cold acclimation-recruited UCP1-dependent thermogenic capacity (Figure 7h *versus* Figure 7f). Despite containing only ∼20 % of the UCP1 present in control mice, cold-acclimated UCP1^Myf5cKO^ mice exhibited a UCP1-dependent thermogenic response indistinguishable from that of control mice (Figure 7h).

We next examined whether thermogenic activation was associated with local tissue hypoxia. Hypoxia was analyzed in both beige and brown adipose tissue from the same mice. Because beige adipose tissue retains UCP1 expression in UCP1^Myf5cKO^ mice and has been proposed as a site of UCP1-independent thermogenesis, the main data are shown for beige adipose tissue (Figure 7i–t), whereas brown adipose tissue is presented in Figure S6h–s. Sections were stained for UCP1, hypoxia adducts, and perilipin to identify thermogenic adipocytes, local hypoxia, and the entire adipocyte population, respectively.

CL-induced thermogenesis was associated with pronounced hypoxic regions in adipose tissues containing UCP1. This pattern was evident in beige adipose tissue from both control and UCP1^Myf5cKO^ mice (Figure 7j,r) as well as in brown adipose tissue from control mice (Figure S6i). In contrast, brown adipose tissue from UCP1^Myf5cKO^ mice, which is largely depleted of UCP1-expressing adipocytes, exhibited only rare and minor hypoxic regions confined close to residual UCP1-positive cells (Figure S6q).

In global UCP1-KO mice, no hypoxia was detected in either beige (Figure 7n) or brown adipose tissue (Figure S6m). This is consistent with the absence of a CL-induced thermogenic response in these mice (Figure 7b, open circles), indicating that oxygen consumption does not exceed the capacity for oxygen delivery under the conditions tested. Importantly, any substantial UCP1-independent thermogenic mechanism would be expected to increase oxygen consumption, induce tissue hypoxia, or both. Neither response was observed, arguing against the existence of physiologically relevant UCP1-independent thermogenesis under these conditions.

Together, these findings indicate that, in cold-acclimated mice, oxygen (and presumably nutrients) availability, rather than UCP1 abundance, constrains maximal thermogenic responses *in vivo*.

## Discussion

The present study demonstrates that in fully cold-acclimated mice, adaptive nonshivering thermogenesis is not constrained by the abundance of UCP1. Selective ablation of UCP1 in thermogenic adipocytes of myogenic origin reduced total UCP1 content by approximately 80 %, yet cold acclimation-recruited thermogenic capacity remained largely preserved. In contrast, complete UCP1 deficiency totally abolished adrenergically induced thermogenesis, demonstrating that although UCP1 is indispensable for adaptive nonshivering thermogenesis, only a fraction of the total UCP1 normally present is required to sustain maximal thermogenic capacity. Instead, our findings indicate that maximal thermogenic output is constrained by the capacity to sustain UCP1-dependent oxidative metabolism, rather than by UCP1 abundance itself.

### Why is UCP1 present in excess?

The preservation of thermogenic capacity despite such a marked reduction in UCP1 abundance indicates that maximal thermogenic output is not determined by the amount of UCP1 present. This conclusion is consistent with our previous finding that UCP1 content in cold-acclimated mice was approximately 40-fold higher than in thermoneutral mice, whereas thermogenic capacity was increased only approximately 4-fold^6^, and with the proposal by Lindsund et al.^35^ that the relationship between UCP1 abundance and thermogenic output is saturable rather than proportional. One explanation for this apparent excess is that UCP1 accumulation reflects prolonged sympathetic stimulation during thermogenic recruitment. Norepinephrine not only mediates recruitment and activates thermogenesis but also promotes *Ucp1* transcription^20^, such that sustained adrenergic stimulation progressively increases UCP1 abundance beyond that required for maximal heat production. Excess UCP1 may nevertheless provide physiological advantages by constituting a functional reserve under physiological conditions that impose greater thermogenic demands than those examined here. This possibility is consistent with the reserve thermogenic capacity observed in cold-acclimated control mice. Whether excess UCP1 also contributes to the spatial organization of thermogenic activity within brown adipose tissue remains an interesting possibility. In support of this idea, hypoxia developed only in discrete regions of thermogenically active adipose tissue during maximal adrenergic stimulation, indicating that thermogenic activity is heterogeneous across the depot.

### What limits thermogenic capacity?

Having established that UCP1 abundance does not constrain maximal thermogenic output *in vivo*, the question becomes what does. At the mitochondrial level, preserved UCP1-dependent respiration despite reduced UCP1 abundance supports the interpretation that fully recruited brown-fat mitochondria contain more UCP1 than is required to sustain maximal UCP1-dependent respiration. This interpretation is consistent with respiratory capacity rather than UCP1 abundance limiting mitochondrial thermogenesis. At the tissue level, pronounced hypoxia during maximal adrenergic stimulation indicates that oxygen consumption can locally exceed oxygen availability despite the marked increases in vascularization and perfusion accompanying cold acclimation. Together, these observations suggest that adaptive thermogenesis is constrained by the capacity to sustain UCP1-dependent oxidative metabolism rather than by the abundance of UCP1 itself.

### Little evidence for a physiologically relevant contribution of proposed UCP1-independent thermogenic pathways

Despite the markedly reduced UCP1 content in UCP1^Myf5cKO^ mice, we found no evidence for compensatory recruitment of proposed UCP1-independent thermogenic pathways^24–32^ at the level of the adipose depot. Likewise, cold acclimation recruited only a small adrenergically inducible thermogenic response in global UCP1-KO mice, consistent with previous reports^13,14,35^. Furthermore, maximal adrenergic stimulation induced pronounced hypoxia only in UCP1-containing adipose tissues, whereas no hypoxia was detected in adipose tissues lacking UCP1. Although differences in tissue perfusion cannot formally be excluded, the absence of hypoxia in UCP1-deficient adipose tissue is most readily explained by its markedly lower oxygen consumption. Together, these findings provide little evidence that proposed UCP1-independent mechanisms make a physiologically relevant contribution to cold acclimation-recruited nonshivering thermogenesis.

### Why does the thermogenic ‘system’ contain multiple adipose depots and adipocyte lineages?

The preservation of thermogenic capacity despite selective loss of UCP1 in the major thermogenic adipocyte lineage may also provide new insight into the organization of the thermogenic adipose system. If maximal thermogenic output is constrained by local oxygen availability rather than UCP1 abundance, the spatial organization of thermogenic adipose tissue may have evolved to distribute thermogenic activity across multiple vascular territories, thereby alleviating local oxygen limitations while supporting high whole-body thermogenic output.

Similarly, the coexistence of multiple thermogenic adipocyte lineages may enhance the robustness of adaptive thermogenesis. Despite the marked reduction in UCP1 within Myf5-derived adipocytes, adaptive thermogenesis remained largely preserved, demonstrating that non-Myf5-derived thermogenic adipocytes can substantially contribute to whole-body thermogenesis when the major thermogenic lineage is compromised. Although selective loss of UCP1 in Myf5-derived adipocytes is not a physiological situation, this model may reveal an intrinsic resilience of the thermogenic adipose system. Whether distinct thermogenic adipocyte lineages also perform specialized physiological functions beyond heat production remains an important question.

### Translational implications

Our findings have important implications for therapeutic strategies aimed at harnessing thermogenic adipose tissue for the treatment of obesity and metabolic disease^8,46–48^. These results suggest that UCP1 abundance should not be considered the only determinant of maximal thermogenic capacity. Rather, UCP1 abundance reflects the degree of thermogenic adipocyte recruitment, whereas maximal thermogenic output depends on the broader metabolic and physiological context supporting UCP1 function, including mitochondrial respiratory capacity and oxygen availability. Consequently, therapeutic strategies focused solely on increasing UCP1 expression may not yield proportional increases in energy expenditure. Instead, successful interventions may require coordinated enhancement of the metabolic competence of thermogenic adipose tissue to fully realize its thermogenic potential.

Collectively, our findings demonstrate that adaptive nonshivering thermogenesis requires UCP1, yet in cold-acclimated mice only a fraction of the UCP1 normally present is required to sustain maximal thermogenic capacity. Rather than being constrained by UCP1 abundance, maximal thermogenic output is ultimately limited by the capacity to sustain UCP1-dependent oxidative metabolism. These findings provide a revised framework for understanding both the physiological regulation of adaptive thermogenesis and its therapeutic exploitation.

## Methods

### Experimental model

All animal experiments were approved by the Animal Ethics Committee of the North Stockholm region.

Mice with selective ablation of UCP1 in thermogenic adipocytes of myogenic origin were generated by crossing mice carrying floxed Ucp1 alleles (Ucp1^fl/fl^) with Myf5-Cre mice (B6.129S4-Myf5^tm3^(cre)^Sor^/J, Jackson Laboratory stock no. 007893), which carry a Cre knock-in allele at the endogenous Myf5 locus and mediate recombination in cells derived from the Myf5 lineage. The resulting UCP1^Myf5cKO^ mice have the genotype Myf5-Cre^+/-^; Ucp1^fl/fl^.

The Ucp1^fl/fl^ mouse line was generated within the framework of the EUCOMM program^49,50^ (see details in Figure S1). The original *Ucp1^tm^*^1a^ allele contained a lacZ reporter cassette, a neomycin resistance (neo) cassette, two FRT sites, and three loxP sites. *Ucp1^tm^*^1a^ mice were crossed with a Flp recombinase-expressing mouse line^51^, resulting in excision of the lacZ and neo cassettes together with one FRT site and one loxP site, thereby generating the conditional *Ucp1^tm^*^1c^ allele (Ucp1^fl^). Subsequent crossing with Myf5-Cre mice produced the *Ucp1^tm^*^1d^ allele (UCP1^Myf5cKO^), in which exon 2 of the *Ucp1* gene is deleted in cells of the Myf5 lineage, a lineage that contributes substantially to brown adipocytes.

ROSA26-loxP-STOP-loxP-EYFP reporter mice (B6.129X1-Gt(ROSA)26Sor^tm1^(EYFP)^Cos^/J, Jackson Laboratory stock no. 006148)^52^, hereafter referred to as Rosa-YFP^+/-^ or Rosa-YFP^+/+^ mice, carry an enhanced yellow fluorescent protein (EYFP) reporter at the ROSA26 locus that is activated following Cre-mediated excision of a loxP-flanked STOP cassette, enabling permanent lineage tracing of Cre-expressing cells and their descendants. Triple-mutant Myf5-Cre^+/-^; Ucp1^fl/fl^; Rosa-YFP^+/-^ mice were generated by crossing Myf5-Cre^+/-^; Ucp1^fl/fl^ mice with Rosa-YFP^+/-^ mice. Offspring carrying the desired alleles were subsequently intercrossed to obtain mice homozygous for the Rosa-YFP reporter allele (Rosa-YFP^+/+^), thereby increasing reporter fluorescence.

Global UCP1-KO mice were the progeny of those described in^12^, backcrossed to the C57Bl/6J background. The mice were bred and maintained in-house as homozygous lines (UCP1-knockout and wild-type). To avoid genetic drift, UCP1-knockout and wild-type lines were regularly intercrossed.

Before the start of the experiment, mice were housed at 22–24 °C in individually ventilated cages containing poplar bedding, nesting material, a paper tube, and a chewing stick, under a 12:12 h light-dark cycle with free access to chow diet (Brogaarden Altromin 1324) and water. At the start of the experiment, mice aged approximately 8 weeks were single-housed and either acclimated to thermoneutrality (30 °C) or, gradually acclimated to cold by housing them at 18 °C for approximately 1 week followed by 4 °C for 6–8 weeks.

At the end of the acclimation period, mice underwent one or more of the following procedures, depending on the experimental cohort: body composition analysis, measurement of resting metabolic rate at the acclimation temperature, assessment of thermogenic capacity using either the CL test or the norepinephrine test in the Promethion system, and/or euthanasia followed by tissue collection.

### Sampling of tissues

At the indicated time points (i.e., at the end of the experiments), mice were euthanized by CO₂ inhalation. Interscapular brown adipose tissue (IBAT), axillary brown adipose tissue (aBAT), cervical brown adipose tissue (cBAT), perirenal brown adipose tissue (prBAT), and inguinal white adipose tissue (ingWAT) were quantitatively dissected. All depots were snap-frozen in liquid nitrogen and stored at −80 °C for molecular analyses. For histological analyses, one side of the IBAT and ingWAT depots was immersion-fixed in formaldehyde solution, whereas the corresponding contralateral depots were used for molecular analyses.

### Protein analysis

#### Sample processing and protein quantification

Frozen tissues were homogenized in a modified RIPA buffer (50 mM Tris·HCl, pH 7.4, 1 % Triton X-100, 150 mM NaCl, 1 mM EDTA) with freshly added 1 mM Na_3_VO_4_, 10 mM NaF and protease inhibitor cocktail (Complete-Mini, Roche, 04693124001) at a specific ratio, typically 1:10 (w/vol). The homogenates, after freezing (in liquid nitrogen) and subsequent defrosting to ensure complete lysis of adipose cells, were centrifuged at 14000 g for 15 min. The top fat layer was discarded, and the lysate (infranatant) was carefully aspirated using a 1 ml syringe and 27 G needle.

The protein concentration in the lysate was determined using the Lowry method^53^. The total protein content in the depot was calculated by multiplying the protein concentration (in μg protein/μl lysate) by the dilution factor and the wet tissue weight.

#### Western blot analysis

An equal volume of reducing sample buffer (125 mM Tris·HCl, pH 6.8, 4 % (wt/vol) SDS, 20 % (vol/vol) glycerol, 100 mM dithiothreitol, and 0.1 % (wt/vol) bromphenol blue) was added to each sample. Equal amounts of protein were separated by SDS-PAGE in high-resolution 12 % polyacrylamide gel (acrylamide/bis-acrylamide = 175/1). Proteins were transferred to polyvinylidene difluoride membranes (BioRad) in 48 mM Tris·HCl, 39 mM glycine, 0.037 (wt/vol) SDS and 15 % (vol/vol) methanol, using a semi-dry electrophoretic transfer cell (Bio-Rad Trans-Blot SD, Bio-Rad) at 1.2 mA/cm^2^ for 90 min. After washing, the membrane was blocked in 5 % milk in Tris-buffered Saline-Tween for 1 h at room temperature and probed with the indicated antibodies overnight at 4 °C. The immunoblot was visualized with appropriate horseradish peroxidase-conjugated secondary antibodies and enhanced chemiluminescence (Clarity Western ECL Substrate, Bio-Rad) in a charge-coupled device camera (ChemiDoc XRS^+^, Bio-Rad). Analysis of the blots was performed using Image Lab 6.1 software (Bio-Rad). Samples loaded on different membranes were compared by normalization of band intensity with a standard sample of brown fat loaded on all membranes.

Antibodies used for Western blotting were: UCP1 (rabbit polyclonal antibody raised against the C-terminal decapeptide, custom-made, C10; 1:15 000), tyrosine hydroxylase (rabbit monoclonal; Abcam, ab137869; 1:2000), OxPhos Rodent WB Antibody Cocktail (Invitrogen, 45-8099; 1:10000), HADHA (Abcam, ab203114; 1:2000), and β-actin (Invitrogen, MA1-140; 1:5000).

Western blot data were not normalized to a housekeeping protein because no protein with stable expression across adipose depots and experimental conditions could be identified. Instead, equal protein loading was used for normalization. Because several target proteins were absent or expressed at very low levels in some samples, successful sample loading and transfer were verified by probing membranes for β-actin.

### Immunohistochemistry

Adipose tissue depots designated for histological analysis were immersion-fixed in 4 % alcoholic formaldehyde (4% formaldehyde in ethanol) for 24 h, dehydrated, and embedded in paraffin according to a standard procedure^54^. Sections (5 μm) were cut using a microtome (Leica RM2255, Leica Microsystems) and mounted on SuperFrost® Plus adhesion slides (VWR International bvba, Leuven, Belgium). Sections were deparaffinized and rehydrated prior to antigen retrieval.

Antigen unmasking was performed by heating sections in citrate buffer (10 mM sodium citrate, pH 6) for 30 min in a water bath, followed by cooling for 30 min at room temperature. To reduce autofluorescence, sections were incubated with 0.3 % Sudan Black B (Sigma-Aldrich, 199664) in 70 % ethanol for 30 min at room temperature in a humid chamber containing ethanol vapour, followed by washing in PBS.

For multiplex immunostaining (essentially as described in^37,55^), sequential staining with tyramide signal amplification (TSA) reagents (Thermo Fisher Scientific) enabled the use of primary antibodies raised in the same host species. Alexa Fluor 488- and Alexa Fluor 594-labelled tyramides were used according to the manufacturer’s instructions (kits B40943 and B40944, respectively). Endogenous peroxidase activity was quenched using 3 % hydrogen peroxide (kit component C2) for 1 h at room temperature. Sections were blocked in 3 % BSA in PBS for 2 h at room temperature prior to antibody incubation. Primary antibodies were diluted in 1 % BSA in PBS and applied at ∼30 μl per section, followed by incubation for 24 h at 4 °C in a humidified chamber.

Nuclei were visualized with Hoechst 33258 (1 μg/ml; Sigma-Aldrich, 861405) for 10 min, followed by washing in PBS and mounting with ProLong Gold Antifade Reagent (Molecular Probes, P36934).

#### Sequential TSA-based immunostaining procedures

UCP1–perilipin staining (Figure 1)

Sections were first incubated overnight with UCP1 antibody (rabbit polyclonal raised against C-terminal decapeptide, custom-made, C10; 1:1000), followed by HRP-conjugated secondary antibody (Component B, B40943 kit; 1 h) and TSA development with Alexa Fluor 488 (10 min). Antibodies were removed by heat-induced antibody stripping in citrate buffer (30 min), followed by cooling and blocking in 3 % BSA (30 min). Sections were then incubated overnight with perilipin antibody (rabbit monoclonal, Cell Signaling Technology, #9349; 1:500) and visualized using Alexa Fluor 594-conjugated goat anti-rabbit secondary antibody (Molecular Probes, A11037; 1:250). Imaging was performed using a Zeiss LSM 780 confocal microscope (Carl Zeiss MicroImaging).

YFP–UCP1 staining (Figure S2)

Sections were incubated overnight with anti-GFP antibody (Abcam, ab6673; 1:750), followed by HRP-conjugated donkey anti-goat secondary antibody (Invitrogen, A15999; 1:500, 1 h) and TSA development with Alexa Fluor 594 (B40944; 10 min). Sections were then incubated overnight with the UCP1 antibody (rabbit polyclonal raised against the C-terminal decapeptide, custom-made, C10; 1:1000) and visualized using Alexa Fluor 488-conjugated chicken anti-rabbit secondary antibody (Molecular Probes, A21470; 1:250, 2 h). Imaging was performed as above.

Tyrosine hydroxylase–perilipin–UCP1 staining (Figure 5)

Sections were sequentially incubated overnight with tyrosine hydroxylase (rabbit monoclonal; Abcam, ab137869; 1:350), perilipin (rabbit monoclonal; Cell Signaling Technology, #9349; 1:1000), and UCP1 (rabbit monoclonal; Abcam, EPR20381; 1:250) antibodies. Tyrosine hydroxylase and perilipin were visualized using TSA (Alexa Fluor 488 and 594, respectively), with heat-induced antibody stripping between steps. UCP1 was detected using Alexa Fluor 647-conjugated goat anti-rabbit secondary antibody (Molecular Probes, A21245; 1:250, 2 h). Imaging was performed using a Zeiss LSM 780 confocal microscope.

UCP1–pimonidazole–perilipin staining (Figures 7 and S6)

Sections were first incubated overnight with UCP1 antibody (rabbit monoclonal; Abcam, EPR20381; 1:1000), followed by HRP-conjugated secondary antibody (Component B, B40944; 1 h) and TSA development with Alexa Fluor 594 (10 min; B40944). Antibodies were removed by heat-induced antibody stripping in citrate buffer (30 min), followed by cooling and blocking with hydrogen peroxide (30 min), 3 % BSA (30 min), and endogenous biotin blocking kit (Molecular Probes, E21390).

Sections were then incubated overnight with biotinylated anti-pimonidazole antibody (mouse IgG1 monoclonal, clone 4.3.11.3; Hypoxyprobe™, 1:75), followed by DyLight™ 488-conjugated streptavidin (Thermo Fisher Scientific; 1:100, 2 h). Subsequently, sections were incubated overnight with perilipin antibody and visualized using Alexa Fluor 647-conjugated goat anti-rabbit secondary antibody (Molecular Probes, A21245; 1:250, 2 h). Imaging was performed using a Zeiss LSM 780 (Figure 7) or Zeiss LSM 800 (Figure S6) confocal microscope.

### mRNA analysis

#### RNA isolation and cDNA synthesis

Frozen tissues were homogenized in TRI Reagent (T9424, Sigma-Aldrich), and the chloroform-isopropanol method was used to isolate RNA according to the Sigma-Aldrich TRI Reagent protocol. The RNA concentrations in the samples were measured with a Thermo Scientific NanoDrop One Spectrophotometer. The High-Capacity cDNA Reverse Transcription Kit (4368814, Applied Biosystems) was used to reverse transcribe 500 ng of total RNA into cDNA in a total volume of 20 μl. After the reaction was completed, cDNA was diluted 10 times in water.

#### Real-time qPCR

All primers were validated before use to ensure good amplification efficiency (90–110%) and specificity (controlled for by melting curve analysis and inclusion of control samples in which the reverse transcriptase had been left out of the reaction). Gene-specific primers (see Table S1) and PowerUp™ SYBR™ Green Master Mix for qPCR (A25742, Applied Biosystems) were premixed in a total volume of 11 μl. The final primer concentration used was 0.3 μM. Two microliters of the diluted cDNA were added to the premixed primer solution to a total volume of 13 μl. All samples were run in triplicate. The Bio-Rad CFX Connect Real-Time system was used to perform the real-time quantitative polymerase chain reaction. The samples were preheated 2 min at 50 °C and 10 min at 95 °C, after which 40 cycles of 15 s at 95 °C and 1 min at 60 °C were run. The real-time qPCR reaction was followed by melting curve analysis.

The ΔCt method was used to calculate relative changes in mRNA abundance. Ct values for 18S rRNA were subtracted from the Ct values of each analyzed gene (ΔCt method) to adjust for variability in cDNA synthesis. These ΔCt values were antilog-transformed (2^−ΔCt^) to determine changes in mRNA abundance. Reference gene expression was analyzed as 2^−Ct^ and was generally similar among samples (Figure S3a).

### Isolation and analysis of brown-fat mitochondria

#### Isolation of brown-fat mitochondria

Cold-acclimated mice (as described above) were anesthetized for approximately 1 min with a mixture of 79 % CO_2_ and 21 % O_2_ and decapitated. Brown fat depots (interscapular and axillary) were placed in ice-cold medium containing 250 mM sucrose, 10 mM TES (pH 7.2), 1 mM EGTA, 0.1 % fatty-acid-free BSA and used for isolation of mitochondria. Preparations of brown-fat mitochondria from control and heterozygous mice were routinely made and examined in parallel.

Brown-fat mitochondria were isolated as described^56^. The tissues were finely minced with scissors and homogenized in a Potter homogenizer with a Teflon pestle. Throughout the isolation process, tissues were kept at 0–2 °C. Mitochondria were isolated by differential centrifugation as follows. The homogenate was filtered through cotton gauze and centrifuged at 8800 g for 10 min at 2 °C, using a Beckman Coulter^TM^ Allegra^TM^ 25R centrifuge. The resulting supernatant, containing floating fat, was discarded. The pellet was resuspended in ice-cold medium 250 mM sucrose, 10 mM TES (pH 7.2), 1 mM EGTA, 0.6 % fatty-acid-free BSA. The resuspended homogenate was centrifuged at 800 g for 10 min, and the resulting supernatant was centrifuged again at 8800 g for 10 min. The resulting mitochondrial pellet was resuspended in 100 mM KCl, 20 mM K^+^-TES (pH 7.2), 1 mM EDTA, and 0.6 % fatty acid-free BSA (Cat#10775835001, Roche) and centrifuged at 8800 g for 10 min at 2 °C. The final mitochondrial pellets were resuspended by hand homogenization in a small glass homogenizer in the same medium to a protein concentration of 30-40 mg/ml. The concentration of mitochondrial protein was measured using Lowry method^53^ with BSA as a standard. The mitochondria were stored on ice prior to respiration measurements.

#### Mitochondrial oxygen consumption

Oxygen consumption was measured in a high-resolution oxygraph (Oroboros O2k-FluoRespirometer, Austria) at 37 °C. The mitochondria (0.125 μg protein) were incubated in a medium consisting of 125 mM sucrose, 20 mM K^+^-TES (pH 7.2), 2 mM MgCl_2_, 1 mM EDTA, 4 mM KH₂PO₄, and 0.1 % fatty acid-free BSA.

The respiratory activity of the mitochondria was measured in the presence of 5 mM malate + 5 mM pyruvate + 0.5 mM octanoyl-L-carnitine. UCP1-dependent thermogenesis was quantified as the respiration that was inhibited by 2 mM GDP. The activity of the phosphorylation system was quantified as the maximal oxygen consumption rate after the addition of 450 μM ADP minus the basal respiration. Basal respiration (“proton leak”) was measured as the residual respiration following the addition of 1.2 μg/ml oligomycin. Maximal oxygen consumption rates were obtained by the addition of 0.6 μM FCCP, followed by 0.2 μM FCCP up to three times.

### Metabolic studies

#### Body composition

Body weight was measured before and after the acclimation period. Body composition (lean and fat mass) was assessed when indicated using magnetic resonance imaging (EchoMRI-700/100 Body Composition Analyzer, Echo Medical Systems, Houston, TX).

#### Resting metabolic rate

Resting metabolic rate was determined by indirect calorimetry at the temperature to which the mice had been acclimated. Oxygen consumption and carbon dioxide production were measured in intact mice using a Promethion indirect calorimetry system (Sable Systems). During the measurements, mice were housed individually in cages identical to their home cages and provided with chow diet and water ad libitum. The cages were maintained under the same light-dark cycle as in the animal facility and contained a small amount of bedding transferred from the home cage. Mice remained in the Promethion system for at least two consecutive nights.

#### Measurement of the nonshivering thermogenic capacity

The capacity for nonshivering thermogenesis was assessed by measuring oxygen consumption in response to either the selective β3-adrenoceptor agonist CL316,243^6,57^ or the sympathomimetic agent norepinephrine (NE)^14^, two established tests of thermogenic capacity.

##### CL316,243 (CL) test

Mice were placed in a Promethion indirect calorimetry system (Sable Systems) at 30 °C overnight. On the following day (approximately 12:00), mice received an intraperitoneal injection of CL (1 mg/kg body weight). Oxygen consumption was subsequently monitored for at least 4.5 h. Mice had ad libitum access to food and water throughout the experiment.

In experiments in which tissue hypoxia was assessed, mice received an intraperitoneal injection of pimonidazole 1.5 h after CL administration and were euthanized 3 h later, resulting in a total CL exposure time of 4.5 h.

Basal metabolic rate was calculated as the average of the 5–10 lowest consecutive stable pre-injection measurements, corresponding to approximately 15–30 min. The CL-induced thermogenic response was calculated as the average oxygen consumption during the final 120 min of the measurement period minus the basal metabolic rate.

##### Norepinephrine (NE) test

The NE test was performed at 30 °C in anesthetized mice to improve measurement reproducibility^21^. Mice were anesthetized with pentobarbital sodium (75–90 mg/kg body weight, intraperitoneally), which, in contrast to volatile anesthetics, does not inhibit brown adipocyte thermogenesis^58^.

Anesthetized mice were placed in 0.7-l glass tubes fitted with a metal grid floor and connected to a Promethion indirect calorimetry system (Sable Systems). Following a baseline measurement period of approximately 15 min, mice were removed from the chambers and injected intraperitoneally with NE (norepinephrine bitartrate; 1 mg/kg body weight for thermoneutral mice and 2 mg/kg body weight for cold-acclimated mice). The mice were then returned to the chambers, and oxygen consumption was monitored for approximately 1 h.

Basal metabolic rate was defined as the average of the 5–9 lowest consecutive stable pre-injection measurements, corresponding to approximately 5–9 min. The NE-induced thermogenic response was calculated as the mean of the three highest post-injection measurements minus the basal metabolic rate.

## Supplementary material

This article contains supplementary material.

## Data availability

All data are contained within the article and supplementary material.

## Acknowledgements

The authors are grateful to Barbara Cannon and Jan Nedergaard for numerous insightful discussions throughout the course of this work. The authors also thank the staff of the Experimental Core Facility for breeding the mice and the Imaging Facility at Stockholm University for assistance with confocal microscopy.

## Author contributions

NP designed the research. QN, CPBS-F and NP performed the experiments. NP, QN, CPBS-F and WP interpreted the data. NP wrote the manuscript, and all authors revised and approved the manuscript.

## References

1. de Jong, J.M., Larsson, O., Cannon, B., and Nedergaard, J. (2015). A stringent validation of mouse adipose tissue identity markers. Am J Physiol Endocrinol Metab 308, E1085–1105. 10.1152/ajpendo.00023.2015.

2. Kalinovich, A.V., de Jong, J.M., Cannon, B., and Nedergaard, J. (2017). UCP1 in adipose tissues: two steps to full browning. Biochimie 134, 127–137. 10.1016/j.biochi.2017.01.007.

3. Shabalina, I.G., Petrovic, N., de Jong, J.M., Kalinovich, A.V., Cannon, B., and Nedergaard, J. (2013). UCP1 in brite/beige adipose tissue mitochondria is functionally thermogenic. Cell Rep 5, 1196–1203. 10.1016/j.celrep.2013.10.044.

4. Rosenwald, M., Perdikari, A., Rulicke, T., and Wolfrum, C. (2013). Bi-directional interconversion of brite and white adipocytes. Nat Cell Biol 15, 659–667. 10.1038/ncb2740.

5. Sanchez-Gurmaches, J., Hung, C.M., and Guertin, D.A. (2016). Emerging Complexities in Adipocyte Origins and Identity. Trends Cell Biol 26, 313–326. 10.1016/j.tcb.2016.01.004.

6. Fischer, A.W., Shabalina, I.G., Mattsson, C.L., Abreu-Vieira, G., Cannon, B., Nedergaard, J., and Petrovic, N. (2017). UCP1 inhibition in Cidea-overexpressing mice is physiologically counteracted by brown adipose tissue hyperrecruitment. Am J Physiol Endocrinol Metab 312, E72–E87. 10.1152/ajpendo.00284.2016.

7. de Jong, J.M.A., Wouters, R.T.F., Boulet, N., Cannon, B., Nedergaard, J., and Petrovic, N. (2017). The beta3-adrenergic receptor is dispensable for browning of adipose tissues. Am J Physiol Endocrinol Metab 312, E508–E518. 10.1152/ajpendo.00437.2016.

8. de Jong, J.M.A., Sun, W., Pires, N.D., Frontini, A., Balaz, M., Jespersen, N.Z., Feizi, A., Petrovic, K., Fischer, A.W., Bokhari, M.H., et al. (2019). Human brown adipose tissue is phenocopied by classical brown adipose tissue in physiologically humanized mice. Nat Metab 1, 830–843. 10.1038/s42255-019-0101-4.

9. Sanchez-Gurmaches, J., and Guertin, D.A. (2014). Adipocytes arise from multiple lineages that are heterogeneously and dynamically distributed. Nat Commun 5, 4099. 10.1038/ncomms5099.

10. Dulloo, A.G., and Miller, D.S. (1984). Energy balance following sympathetic denervation of brown adipose tissue. Can J Physiol Pharmacol 62, 235–240. 10.1139/y84-035.

11. Rothwell, N.J., and Stock, M.J. (1989). Surgical removal of brown fat results in rapid and complete compensation by other depots. Am J Physiol 257, R253–258. 10.1152/ajpregu.1989.257.2.R253.

12. Enerback, S., Jacobsson, A., Simpson, E.M., Guerra, C., Yamashita, H., Harper, M.E., and Kozak, L.P. (1997). Mice lacking mitochondrial uncoupling protein are cold-sensitive but not obese. Nature 387, 90–94. 10.1038/387090a0.

13. Golozoubova, V., Hohtola, E., Matthias, A., Jacobsson, A., Cannon, B., and Nedergaard, J. (2001). Only UCP1 can mediate adaptive nonshivering thermogenesis in the cold. FASEB J 15, 2048–2050. 10.1096/fj.00-0536fje.

14. Golozoubova, V., Cannon, B., and Nedergaard, J. (2006). UCP1 is essential for adaptive adrenergic nonshivering thermogenesis. Am J Physiol Endocrinol Metab 291, E350–357. 10.1152/ajpendo.00387.2005.

15. Challa, T.D., Dapito, D.H., Kulenkampff, E., Kiehlmann, E., Moser, C., Straub, L., Sun, W., and Wolfrum, C. (2020). A Genetic Model to Study the Contribution of Brown and Brite Adipocytes to Metabolism. Cell Rep 30, 3424–3433 e3424. 10.1016/j.celrep.2020.02.055.

16. Timmons, J.A., Wennmalm, K., Larsson, O., Walden, T.B., Lassmann, T., Petrovic, N., Hamilton, D.L., Gimeno, R.E., Wahlestedt, C., Baar, K., et al. (2007). Myogenic gene expression signature establishes that brown and white adipocytes originate from distinct cell lineages. Proc Natl Acad Sci U S A 104, 4401–4406. 10.1073/pnas.0610615104.

17. Seale, P., Bjork, B., Yang, W., Kajimura, S., Chin, S., Kuang, S., Scime, A., Devarakonda, S., Conroe, H.M., Erdjument-Bromage, H., et al. (2008). PRDM16 controls a brown fat/skeletal muscle switch. Nature 454, 961–967.

18. Petrovic, N., Walden, T.B., Shabalina, I.G., Timmons, J.A., Cannon, B., and Nedergaard, J. (2010). Chronic peroxisome proliferator-activated receptor gamma (PPARgamma) activation of epididymally derived white adipocyte cultures reveals a population of thermogenically competent, UCP1-containing adipocytes molecularly distinct from classic brown adipocytes. J Biol Chem 285, 7153–7164. 10.1074/jbc.M109.053942.

19. Shamsi, F., Piper, M., Ho, L.L., Huang, T.L., Gupta, A., Streets, A., Lynes, M.D., and Tseng, Y.H. (2021). Vascular smooth muscle-derived Trpv1(+) progenitors are a source of cold-induced thermogenic adipocytes. Nat Metab 3, 485–495. 10.1038/s42255-021-00373-z.

20. Cannon, B., and Nedergaard, J. (2004). Brown adipose tissue: function and physiological significance. Physiol Rev 84, 277–359. 10.1152/physrev.00015.2003.

21. Cannon, B., and Nedergaard, J. (2011). Nonshivering thermogenesis and its adequate measurement in metabolic studies. J Exp Biol 214, 242–253. 10.1242/jeb.050989.

22. Virtue, S., and Vidal-Puig, A. (2013). Assessment of brown adipose tissue function. Front Physiol 4, 128. 10.3389/fphys.2013.00128.

23. Walden, T.B., Hansen, I.R., Timmons, J.A., Cannon, B., and Nedergaard, J. (2012). Recruited vs. nonrecruited molecular signatures of brown, “brite,” and white adipose tissues. Am J Physiol Endocrinol Metab 302, E19–31. 10.1152/ajpendo.00249.2011.

24. Kazak, L., Chouchani, E.T., Jedrychowski, M.P., Erickson, B.K., Shinoda, K., Cohen, P., Vetrivelan, R., Lu, G.Z., Laznik-Bogoslavski, D., Hasenfuss, S.C., et al. (2015). A creatine-driven substrate cycle enhances energy expenditure and thermogenesis in beige fat. Cell 163, 643–655. 10.1016/j.cell.2015.09.035.

25. Rahbani, J.F., Roesler, A., Hussain, M.F., Samborska, B., Dykstra, C.B., Tsai, L., Jedrychowski, M.P., Vergnes, L., Reue, K., Spiegelman, B.M., and Kazak, L. (2021). Creatine kinase B controls futile creatine cycling in thermogenic fat. Nature 590, 480–485. 10.1038/s41586-021-03221-y.

26. Sun, Y., Rahbani, J.F., Jedrychowski, M.P., Riley, C.L., Vidoni, S., Bogoslavski, D., Hu, B., Dumesic, P.A., Zeng, X., Wang, A.B., et al. (2021). Mitochondrial TNAP controls thermogenesis by hydrolysis of phosphocreatine. Nature 593, 580–585. 10.1038/s41586-021-03533-z.

27. Ikeda, K., Kang, Q., Yoneshiro, T., Camporez, J.P., Maki, H., Homma, M., Shinoda, K., Chen, Y., Lu, X., Maretich, P., et al. (2017). UCP1-independent signaling involving SERCA2b-mediated calcium cycling regulates beige fat thermogenesis and systemic glucose homeostasis. Nat Med 23, 1454–1465. 10.1038/nm.4429.

28. Guan, H.P., Li, Y., Jensen, M.V., Newgard, C.B., Steppan, C.M., and Lazar, M.A. (2002). A futile metabolic cycle activated in adipocytes by antidiabetic agents. Nat Med 8, 1122–1128. 10.1038/nm780.

29. Granneman, J.G., Burnazi, M., Zhu, Z., and Schwamb, L.A. (2003). White adipose tissue contributes to UCP1-independent thermogenesis. Am J Physiol Endocrinol Metab 285, E1230–1236. 10.1152/ajpendo.00197.2003.

30. Oeckl, J., Janovska, P., Adamcova, K., Bardova, K., Brunner, S., Dieckmann, S., Ecker, J., Fromme, T., Funda, J., Gantert, T., et al. (2022). Loss of UCP1 function augments recruitment of futile lipid cycling for thermogenesis in murine brown fat. Mol Metab 61, 101499. 10.1016/j.molmet.2022.101499.

31. Bertholet, A.M., Chouchani, E.T., Kazak, L., Angelin, A., Fedorenko, A., Long, J.Z., Vidoni, S., Garrity, R., Cho, J., Terada, N., et al. (2019). H(+) transport is an integral function of the mitochondrial ADP/ATP carrier. Nature 571, 515–520. 10.1038/s41586-019-1400-3.

32. Long, J.Z., Svensson, K.J., Bateman, L.A., Lin, H., Kamenecka, T., Lokurkar, I.A., Lou, J., Rao, R.R., Chang, M.R., Jedrychowski, M.P., et al. (2016). The Secreted Enzyme PM20D1 Regulates Lipidated Amino Acid Uncouplers of Mitochondria. Cell 166, 424–435. 10.1016/j.cell.2016.05.071.

33. Ukropec, J., Anunciado, R.P., Ravussin, Y., Hulver, M.W., and Kozak, L.P. (2006). UCP1-independent thermogenesis in white adipose tissue of cold-acclimated Ucp1-/- mice. J Biol Chem 281, 31894–31908. 10.1074/jbc.M606114200.

34. Keipert, S., Kutschke, M., Ost, M., Schwarzmayr, T., van Schothorst, E.M., Lamp, D., Brachthauser, L., Hamp, I., Mazibuko, S.E., Hartwig, S., et al. (2017). Long-Term Cold Adaptation Does Not Require FGF21 or UCP1. Cell Metab 26, 437–446 e435. 10.1016/j.cmet.2017.07.016.

35. Lindsund, E., Cannon, B., Bengtsson, T., and Nedergaard, J. (2025). Adrenergic stimulation as a means to determine in vivo brown adipose tissue thermogenic capacity. Canadian Journal of Zoology 103, 1–16. 10.1139/cjz-2024-0156.

36. Dieckmann, S., Strohmeyer, A., Willershauser, M., Maurer, S.F., Wurst, W., Marschall, S., de Angelis, M.H., Kuhn, R., Worthmann, A., Fuh, M.M., et al. (2022). Susceptibility to diet-induced obesity at thermoneutral conditions is independent of UCP1. Am J Physiol Endocrinol Metab 322, E85–E100. 10.1152/ajpendo.00278.2021.

37. Naren, Q., Lindsund, E., Bokhari, M.H., Pang, W., and Petrovic, N. (2024). Differential responses to UCP1 ablation in classical brown versus beige fat, despite a parallel increase in sympathetic innervation. J Biol Chem 300, 105760. 10.1016/j.jbc.2024.105760.

38. Shabalina, I.G., Ost, M., Petrovic, N., Vrbacky, M., Nedergaard, J., and Cannon, B. (2010). Uncoupling protein-1 is not leaky. Biochim Biophys Acta 1797, 773–784.

39. Matthias, A., Jacobsson, A., Cannon, B., and Nedergaard, J. (1999). The bioenergetics of brown fat mitochondria from UCP1-ablated mice. Ucp1 is not involved in fatty acid-induced de-energization (“uncoupling”). J Biol Chem 274, 28150–28160. 10.1074/jbc.274.40.28150.

40. Monemdjou, S., Kozak, L.P., and Harper, M.E. (1999). Mitochondrial proton leak in brown adipose tissue mitochondria of Ucp1-deficient mice is GDP insensitive. Am J Physiol 276, E1073–1082. 10.1152/ajpendo.1999.276.6.E1073.

41. Hofmann, W.E., Liu, X., Bearden, C.M., Harper, M.E., and Kozak, L.P. (2001). Effects of genetic background on thermoregulation and fatty acid-induced uncoupling of mitochondria in UCP1-deficient mice. J Biol Chem 276, 12460–12465. 10.1074/jbc.M100466200.

42. Shabalina, I.G., Jacobsson, A., Cannon, B., and Nedergaard, J. (2004). Native UCP1 displays simple competitive kinetics between the regulators purine nucleotides and fatty acids. J Biol Chem 279, 38236–38248. 10.1074/jbc.M402375200.

43. Abreu-Vieira, G., Hagberg, C.E., Spalding, K.L., Cannon, B., and Nedergaard, J. (2015). Adrenergically stimulated blood flow in brown adipose tissue is not dependent on thermogenesis. Am J Physiol Endocrinol Metab 308, E822–829. 10.1152/ajpendo.00494.2014.

44. Foster, D.O., and Frydman, M.L. (1978). Nonshivering thermogenesis in the rat. II. Measurements of blood flow with microspheres point to brown adipose tissue as the dominant site of the calorigenesis induced by noradrenaline. Can J Physiol Pharmacol 56, 110–122. 10.1139/y78-015.

45. Xue, Y., Petrovic, N., Cao, R., Larsson, O., Lim, S., Chen, S., Feldmann, H.M., Liang, Z., Zhu, Z., Nedergaard, J., et al. (2009). Hypoxia-independent angiogenesis in adipose tissues during cold acclimation. Cell Metab 9, 99–109. 10.1016/j.cmet.2008.11.009.

46. Naja Z. Jespersen, A.F., Eline S. Andersen, Eline S. Andersen, Helle B. Hattel, Søren Daugaard, Lone Peijs, Per Bagi, Bo Feldt-Rasmussen, Heidi S. Schultz, Ninna S. Hansen, Rikke Krogh-Madsen, Bente K. Pedersen, Natasa Petrovic, Søren Nielsen, Camilla Scheele (2019). Heterogeneity in the perirenal region of humans suggests presence of dormant brown adipose tissue that contains brown fat precursor cells. Molecular metabolism.

47. Becher, T., Palanisamy, S., Kramer, D.J., Eljalby, M., Marx, S.J., Wibmer, A.G., Butler, S.D., Jiang, C.S., Vaughan, R., Schoder, H., et al. (2021). Brown adipose tissue is associated with cardiometabolic health. Nat Med 27, 58–65. 10.1038/s41591-020-1126-7.

48. Cypess, A.M., Cannon, B., Nedergaard, J., Kazak, L., Chang, D.C., Krakoff, J., Tseng, Y.H., Scheele, C., Boucher, J., Petrovic, N., et al. (2025). Emerging debates and resolutions in brown adipose tissue research. Cell Metab 37, 12–33. 10.1016/j.cmet.2024.11.002.

49. Pettitt, S.J., Liang, Q., Rairdan, X.Y., Moran, J.L., Prosser, H.M., Beier, D.R., Lloyd, K.C., Bradley, A., and Skarnes, W.C. (2009). Agouti C57BL/6N embryonic stem cells for mouse genetic resources. Nat Methods 6, 493–495. 10.1038/nmeth.1342.

50. Skarnes, W.C., Rosen, B., West, A.P., Koutsourakis, M., Bushell, W., Iyer, V., Mujica, A.O., Thomas, M., Harrow, J., Cox, T., et al. (2011). A conditional knockout resource for the genome-wide study of mouse gene function. Nature 474, 337–342. 10.1038/nature10163.

51. Farley, F.W., Soriano, P., Steffen, L.S., and Dymecki, S.M. (2000). Widespread recombinase expression using FLPeR (flipper) mice. Genesis 28, 106–110.

52. Srinivas, S., Watanabe, T., Lin, C.S., William, C.M., Tanabe, Y., Jessell, T.M., and Costantini, F. (2001). Cre reporter strains produced by targeted insertion of EYFP and ECFP into the ROSA26 locus. BMC Dev Biol 1, 4. 10.1186/1471-213x-1-4.

53. Lowry, O.H., Rosebrough, N.J., Farr, A.L., and Randall, R.J. (1951). Protein measurement with the Folin phenol reagent. J Biol Chem 193, 265–275.

54. Cinti, S., Zingaretti, M.C., Cancello, R., Ceresi, E., and Ferrara, P. (2001). Morphologic techniques for the study of brown adipose tissue and white adipose tissue. Methods Mol Biol 155, 21–51. 10.1385/1-59259-231-7:021.

55. Sousa-Filho, C.P.B., and Petrovic, N. (2025). No UCP1 in the kidney. Mol Metab 95, 102127. 10.1016/j.molmet.2025.102127.

56. Cannon, B., and Nedergaard, J. (2008). Studies of thermogenesis and mitochondrial function in adipose tissues. Methods Mol Biol 456, 109–121. 10.1007/978-1-59745-245-8_8.

57. Goldgof, M., Xiao, C., Chanturiya, T., Jou, W., Gavrilova, O., and Reitman, M.L. (2014). The chemical uncoupler 2,4-dinitrophenol (DNP) protects against diet-induced obesity and improves energy homeostasis in mice at thermoneutrality. J Biol Chem 289, 19341–19350. 10.1074/jbc.M114.568204.

58. Ohlson, K.B., Shabalina, I.G., Lennstrom, K., Backlund, E.C., Mohell, N., Bronnikov, G.E., Lindahl, S.G., Cannon, B., and Nedergaard, J. (2004). Inhibitory effects of halothane on the thermogenic pathway in brown adipocytes: localization to adenylyl cyclase and mitochondrial fatty acid oxidation. Biochem Pharmacol 68, 463–477. 10.1016/j.bcp.2004.03.028.

