## Supplementary Figures and Table for "Only a fraction of UCP1 is required to sustain adaptive nonshivering thermogenesis in the cold"

Figure S1

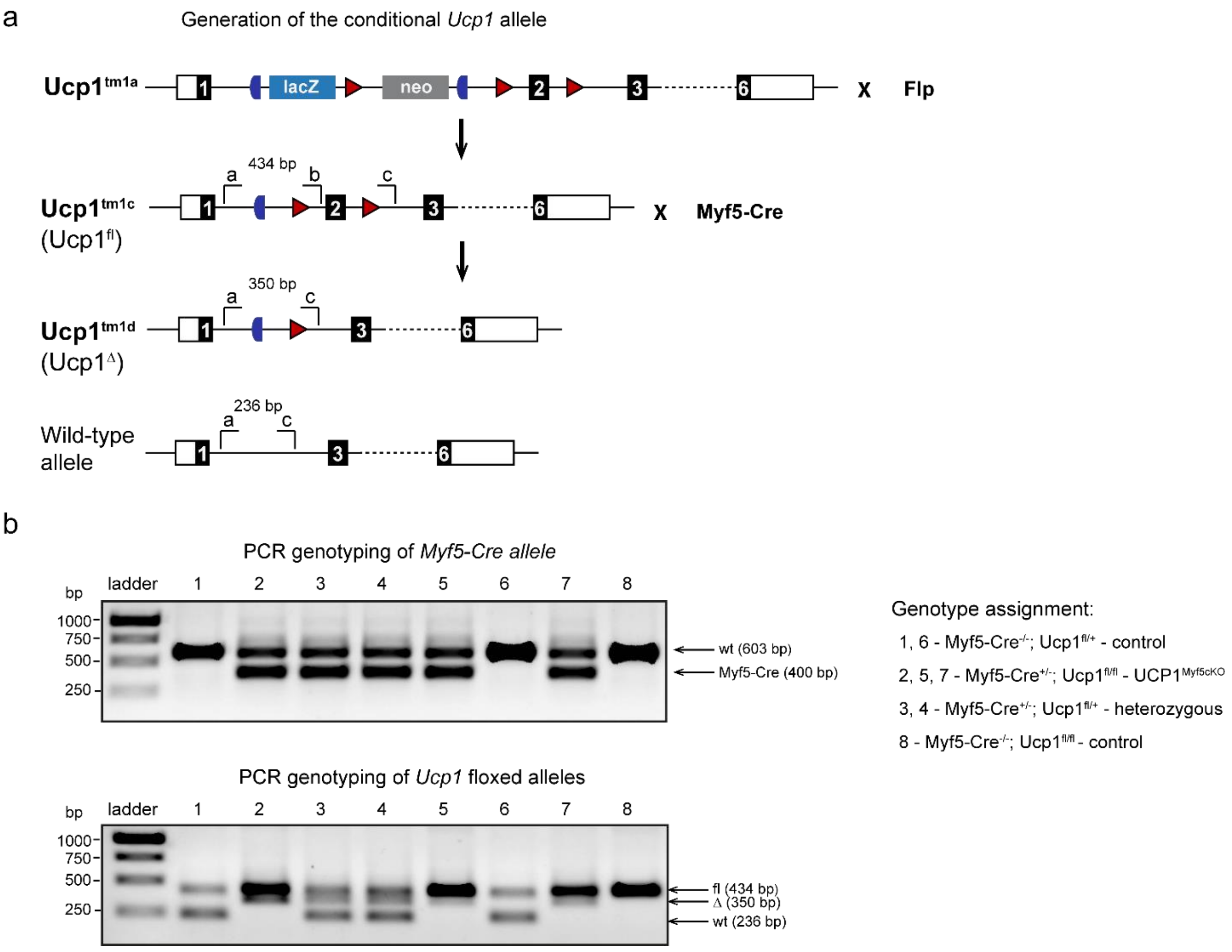


**Figure S1 - Related to Figure 1.**

**Generation of the conditional *Ucp1* allele and representative genotyping.**

**a,** Schematic of the generation of the conditional *Ucp1* allele from the original EUCOMM $Ucp1^{tm1a}$ allele by Flp recombinase-mediated excision of the gene-trap cassette, yielding the conditional-ready floxed $Ucp1^{tm1c}$ ($Ucp1^{fl}$) allele. Cre-mediated recombination deletes exon 2, generating the recombined $Ucp1^{tm1d}$ ($Ucp1^{\Delta}$) allele. Exons are shown as numbered boxes, loxP sites as red triangles, and FRT sites as blue symbols. Lower-case letters (a–c) indicate the binding sites of the primers used for Ucp1 genotyping:
a, AGAACCGCTGTTGATGGGTT; b, TACAATGCAGGCTCCAAACAC;
c, TGTTTGAACTGATGGCGAGC.
The Myf5-Cre allele was genotyped using the following primers:
oIMR7659, CGTAGACGCCTGAAGAAGGTCAACCA;
oIMR7660, ACATTAGAAAACCTGCCAACACC; oIMR7919, ACGAAGTTATTAGGTCCCTCGAC.
**b,** Representative PCR genotyping of the Myf5-Cre allele (upper panel) and Ucp1 alleles (lower panel). The complete genotypes of the mice were: lanes 1 and 6, Myf5-$Cre^{-/-}$; $Ucp1^{fl/+}$ (control mice); lanes 2, 5, and 7, Myf5-$Cre^{+/-}$; $Ucp1^{fl/fl}$ ($UCP1^{Myf5cKO}$ mice); lanes 3 and 4, Myf5-$Cre^{+/-}$; $Ucp1^{fl/+}$ (heterozygous $UCP1^{Myf5cKO}$ mice); lane 8, Myf5-$Cre^{-/-}$; $Ucp1^{fl/fl}$ (control mice). The recombined ($Ucp1^{\Delta}$) allele is detected in DNA from Myf5-Cre-positive mice because ear biopsies contain Myf5-derived dermal cells in which Cre-mediated recombination has occurred.

Figure S2

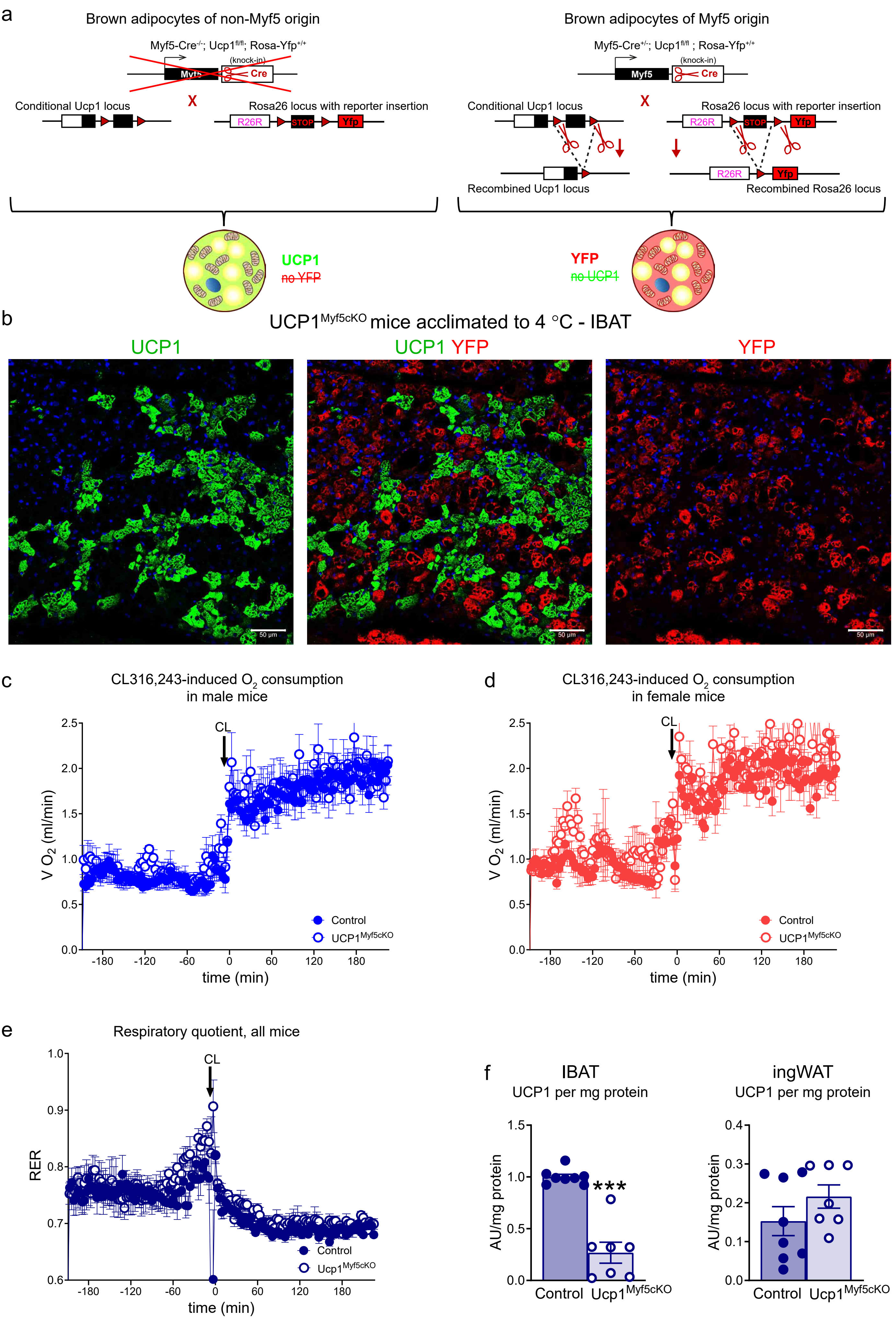

a
Brown adipocytes of non-Myf5 origin
Myf5-Cre-/-; Ucp1fl/fl; Rosa-Yfp+/+
(knock-in)
Myf5
Cre
X
Conditional Ucp1 locus
Rosa26 locus with reporter insertion
R26R
STOP
Yfp
UCP1
no YFP
Brown adipocytes of Myf5 origin
Myf5-Cre+/-; Ucp1fl/fl ; Rosa-Yfp+/+
Recombined Ucp1 locus
Recombined Rosa26 locus
YFP
no UCP1
b
UCP1Myf5cKO mice acclimated to 4 °C - IBAT
UCP1
UCP1 YFP
YFP
50 μm
c
CL316,243-induced O2 consumption in male mice
d
CL316,243-induced O2 consumption in female mice
CL
V O2 (ml/min)
time (min)
Control
UCP1Myf5cKO
e
Respiratory quotient, all mice
RER
Ucp1Myf5cKO
f
IBAT
UCP1 per mg protein
ingWAT
UCP1 per mg protein
AU/mg protein
***
Control

**Figure S2 - Related to Figure 1.**

**Validation of adipocyte lineages and additional characterization of CL-induced thermogenesis.**

**a,** Genetic strategy used to lineage-trace UCP1-positive and UCP1-negative adipocytes within IBAT of $UCP1^{Myf5cKO}$ mice.

**b,** Immunohistochemical analysis of adipose tissue from adult triple-mutant $Myf5\text{-}Cre^{+/-}$; $Ucp1^{fl/fl}$; $Rosa\text{-}Yfp^{+/+}$ mice. Tissues were stained for UCP1 (green), perilipin (red), and nuclei (blue). Scale bar, 50 μm. UCP1 and YFP expression were mutually exclusive, indicating recombination in embryonic Myf5-Cre-expressing cells (UCP1-negative and YFP-positive) but not in precursors of non-Myf5 origin (UCP1-positive and YFP-negative), and confirming that UCP1-positive adipocytes persisting in $UCP1^{Myf5cKO}$ mice are of non-Myf5 origin.

**c–d,** CL316,243 (CL)-induced oxygen consumption in cold-acclimated female mice (c; control, n = 3; $UCP1^{Myf5cKO}$, n = 2) and male mice (d; control, n = 6; $UCP1^{Myf5cKO}$, n = 5).

**e,** Respiratory exchange ratio (RER) in control (n = 8) and $UCP1^{Myf5cKO}$ mice (n = 7) following CL316,243 injection.

**f,** UCP1 protein levels (quantification of immunoblots shown in Figure 1n,p) in IBAT (Figure 1n) and ingWAT (Figure 1p). The mean value for IBAT of control mice was set to 1.0, and all values are expressed relative to this value. Each symbol represents one mouse. Values are means ± SEM. Where not visible, error bars are smaller than the symbols. Asterisks indicate significant differences between control and $UCP1^{Myf5cKO}$ mice (***$P < 0.001$; two-tailed unpaired Student's t-test).

Figure S3

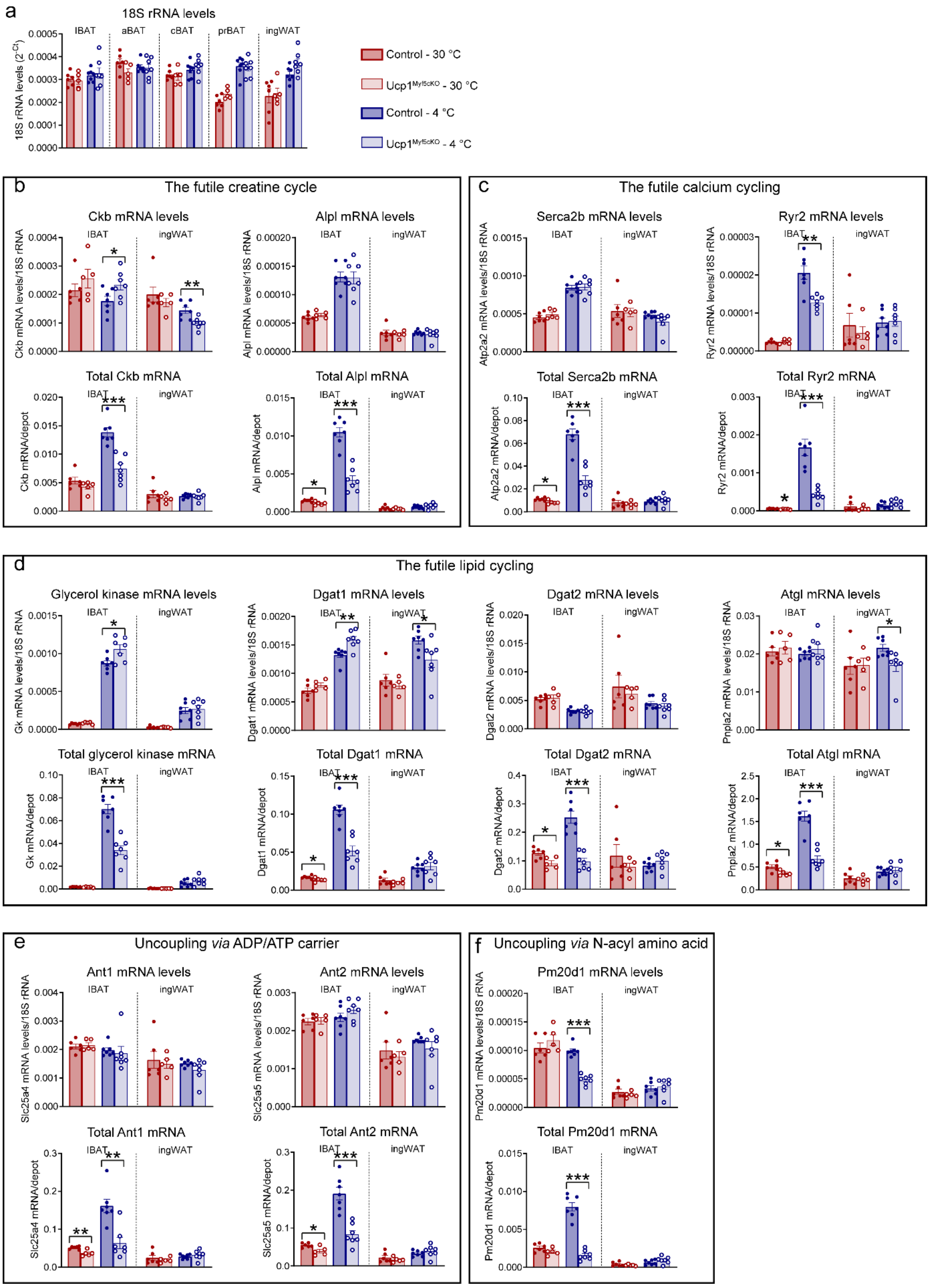
a
18S rRNA levels
IBAT
aBAT
cBAT
prBAT
ingWAT
18S rRNA levels (2^-Ct)
Control - 30 °C
Ucp1Myf5cKO - 30 °C
Control - 4 °C
Ucp1Myf5cKO - 4 °C
b
The futile creatine cycle
Ckb mRNA levels
Ckb mRNA levels/18S rRNA
Alpl mRNA levels
Alpl mRNA levels/18S rRNA
Total Ckb mRNA
Ckb mRNA/depot
Total Alpl mRNA
Alpl mRNA/depot
c
The futile calcium cycling
Serca2b mRNA levels
Atp2a2 mRNA levels/18S rRNA
Ryr2 mRNA levels
Ryr2 mRNA levels/18S rRNA
Total Serca2b mRNA
Atp2a2 mRNA/depot
Total Ryr2 mRNA
Ryr2 mRNA/depot
d
The futile lipid cycling
Glycerol kinase mRNA levels
Gk mRNA levels/18S rRNA
Dgat1 mRNA levels
Dgat1 mRNA levels/18S rRNA
Dgat2 mRNA levels
Dgat2 mRNA levels/18S rRNA
Atgl mRNA levels
Pnpla2 mRNA levels/18S rRNA
Total glycerol kinase mRNA
Gk mRNA/depot
Total Dgat1 mRNA
Dgat1 mRNA/depot
Total Dgat2 mRNA
Dgat2 mRNA/depot
Total Atgl mRNA
Pnpla2 mRNA/depot
e
Uncoupling via ADP/ATP carrier
Ant1 mRNA levels
Slc25a4 mRNA levels/18S rRNA
Ant2 mRNA levels
Slc25a5 mRNA levels/18S rRNA
Total Ant1 mRNA
Slc25a4 mRNA/depot
Total Ant2 mRNA
Slc25a5 mRNA/depot
f
Uncoupling via N-acyl amino acid
Pm20d1 mRNA levels
Pm20d1 mRNA levels/18S rRNA
Total Pm20d1 mRNA
Pm20d1 mRNA/depot

**Figure S3, related to Figure 2**

**Expression of key components of proposed UCP1-independent thermogenic pathways in IBAT and ingWAT of control and UCP1$^{Myf5cKO}$ mice acclimated to thermoneutrality or cold.**

The same IBAT and ingWAT samples as those shown in Figure 2 were analyzed. Expression levels of genes associated with the proposed UCP1-independent thermogenic pathways were normalized to 18S rRNA and are shown in the upper panels for each pathway. Transcript content per whole IBAT and ingWAT depots is shown in the corresponding lower panels. It was calculated by multiplying normalized gene expression levels by the total RNA content of each depot (Figure 2d). **a,** 18S rRNA. **b,** Enzymes of the futile creatine cycle: Ckb and Alpl. **c,** Enzymes involved in futile calcium cycling: Serca2b and Ryr2. **d,** Enzymes of the futile lipid cycle: glycerol kinase (Gk), Dgat1, Dgat2, and Atgl. **e,** Ant1 and Ant2, mediators of ADP/ATP carrier-dependent uncoupling. **f,** Pm20d1, implicated in N-acyl amino acid-mediated uncoupling.

Each symbol represents one mouse. Values are means ± SEM. Asterisks indicate significant differences between control and UCP1$^{Myf5cKO}$ mice (*$P < 0.05$, **$P < 0.01$, ***$P < 0.001$; two-tailed unpaired Student's t-test).

Figure S4

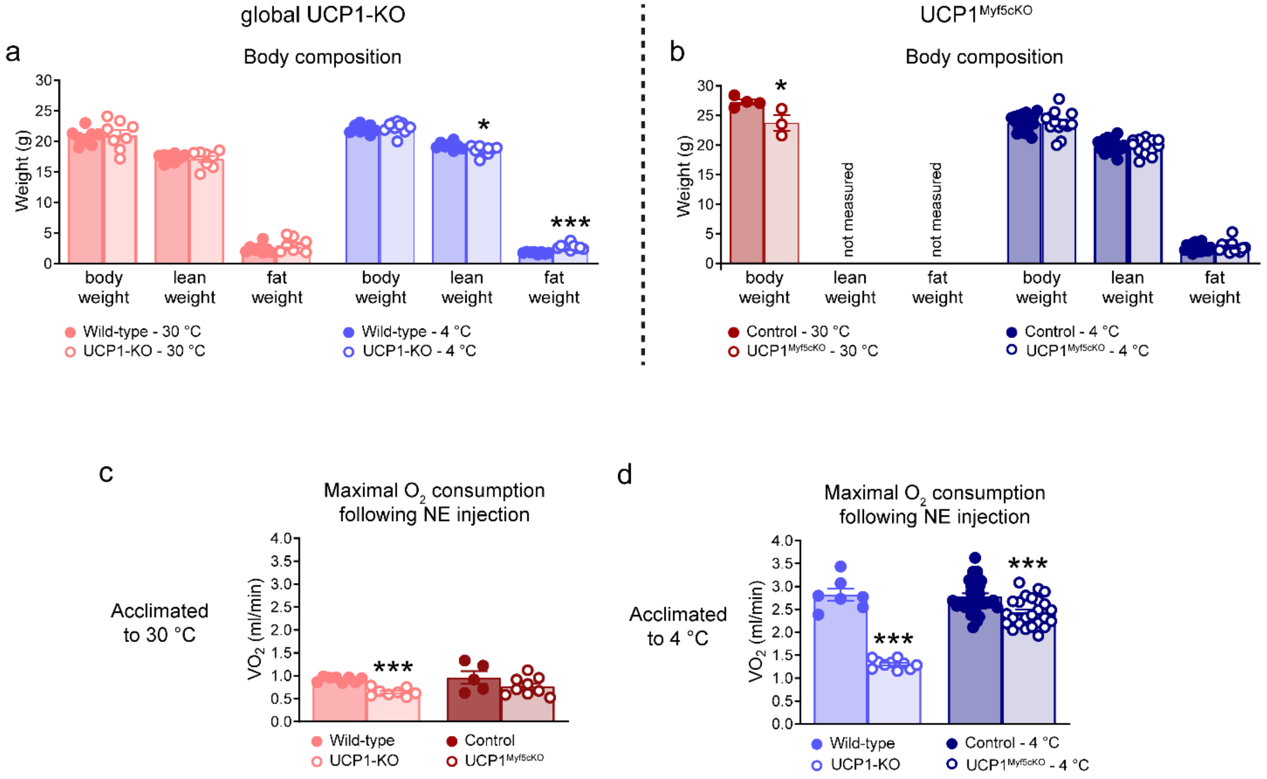


**Figure S4, related to Figure 3**

**Body composition and maximal oxygen consumption of UCP1-KO and UCP1$^{Myf5cKO}$ mice.**

**a,b,** Body composition of the mice presented in Figure 3a–d. **a,** Wild-type mice (30 °C, n = 8; 4 °C, n = 8) and UCP1-KO mice (30 °C, n = 8; 4 °C, n = 10). **b,** Control and UCP1$^{Myf5cKO}$ mice acclimated to 4 °C (control, n = 19; UCP1$^{Myf5cKO}$, n = 13). Body composition was not determined in the corresponding thermoneutral groups.

**c,d,** Maximal oxygen consumption following NE injection in mice acclimated to 30 °C (**c**) or 4 °C (**d**), calculated from the data presented in Figures 3e–h.

Each symbol represents one mouse. Values are means ± SEM. Asterisks indicate significant differences between genotypes (wild-type vs. UCP1-KO; control vs. UCP1$^{Myf5cKO}$; P < 0.05; *** P < 0.001; two-tailed unpaired Student's t-test).

Figure S5

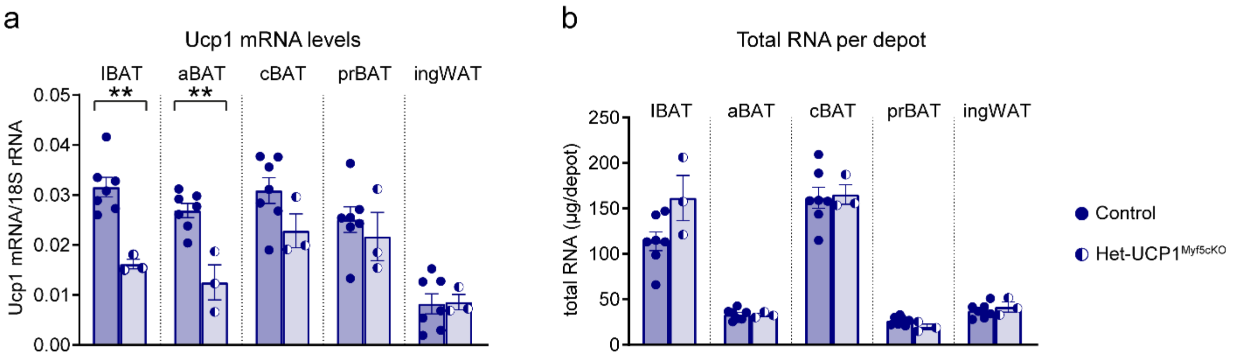


**Figure S5, related to Figure 4**

**Ucp1 mRNA expression and total RNA content across adipose depots in cold-acclimated control and heterozygous Het-UCP1$^{Myf5cKO}$ mice.**

**a,b,** Control (n = 7) and Het-UCP1$^{Myf5cKO}$ (n = 3) mice presented in Figure 4. Interscapular brown adipose tissue (IBAT), axillary BAT (aBAT), cervical BAT (cBAT), perirenal BAT (prBAT), and inguinal white adipose tissue (ingWAT) were analyzed. **a,** Ucp1 mRNA levels normalized to 18S rRNA. **b,** Total RNA content of each depot.

Each symbol represents one mouse. Values are means ± SEM. Asterisks indicate significant differences between control and Het-UCP1$^{Myf5cKO}$ mice **P < 0.01 (two-tailed unpaired Student's t-test).

### Figure S6

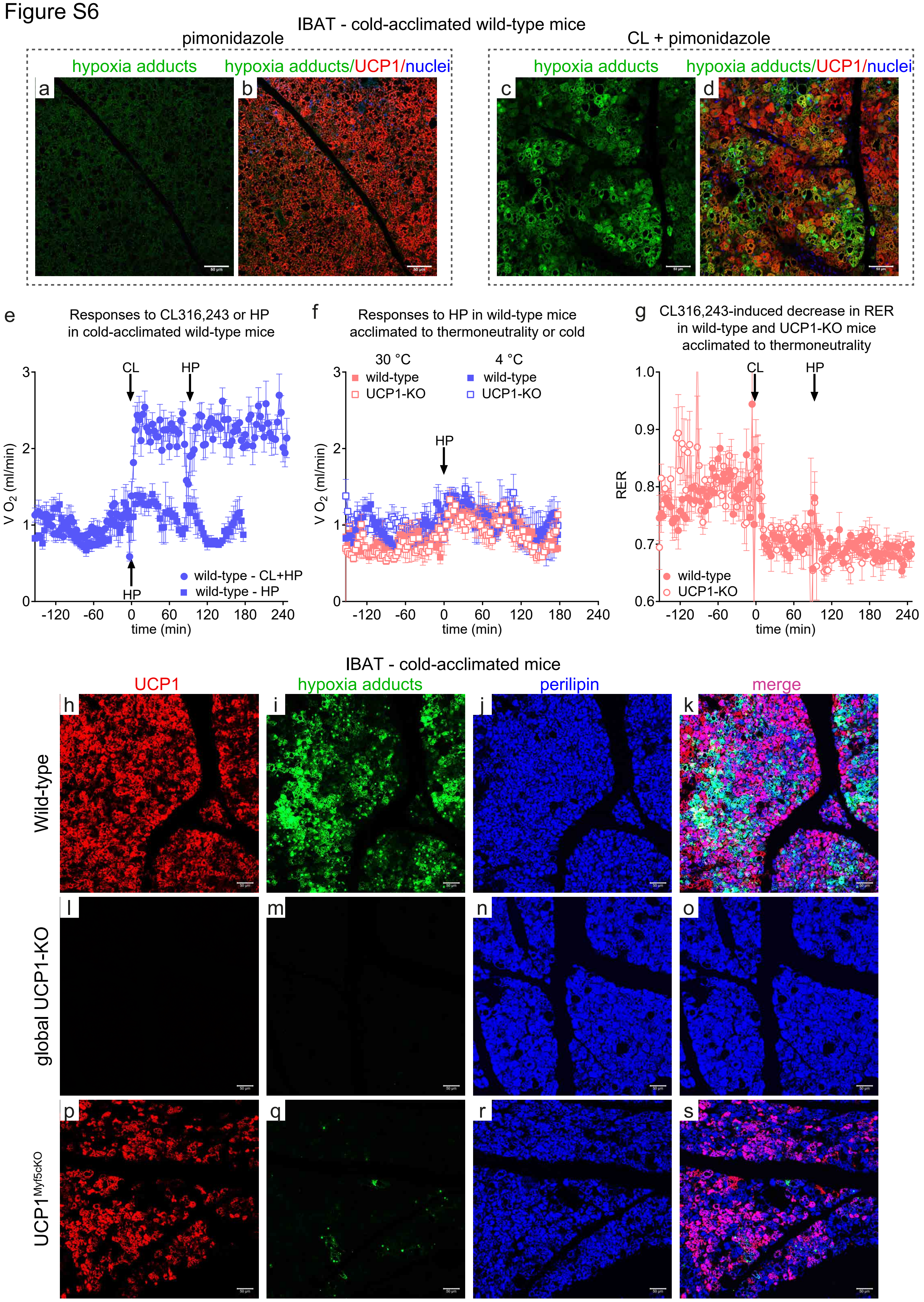
IBAT - cold-acclimated wild-type mice
pimonidazole
CL + pimonidazole
hypoxia adducts
hypoxia adducts/UCP1/nuclei
a
b
c
d
e
Responses to CL316,243 or HP in cold-acclimated wild-type mice
CL
HP
V O2 (ml/min)
wild-type - CL+HP
wild-type - HP
time (min)
f
Responses to HP in wild-type mice acclimated to thermoneutrality or cold
30 °C
4 °C
wild-type
UCP1-KO
g
CL316,243-induced decrease in RER in wild-type and UCP1-KO mice acclimated to thermoneutrality
RER
IBAT - cold-acclimated mice
UCP1
hypoxia adducts
perilipin
merge
Wild-type
global UCP1-KO
UCP1 Myf5cKO
h
i
j
k
l
m
n
o
p
q
r
s

**Figure S6, related to Figure 7. Additional analyses.**

**a–d, Validation of hypoxia detection in IBAT from wild-type mice acclimated to 4 °C. a,b,** Representative confocal images from mice treated with pimonidazole alone. **c,d,** Representative confocal images demonstrating CL-dependent induction of hypoxia adduct staining in IBAT from cold-acclimated wild-type mice treated with CL followed by pimonidazole. Tissues were stained for UCP1 (red), hypoxia adducts (green), and nuclei (blue). For clarity, panels a and c show hypoxia adduct staining only, whereas panels b and d show merged images. Scale bar, 50 μm.

**e, Assessment of injection stress-induced changes in oxygen consumption.** Wild-type mice acclimated to 4 °C were injected with CL followed by pimonidazole (HP; blue circles) or with pimonidazole alone (blue squares). Pimonidazole alone did not elicit an appreciable increase in oxygen consumption. The CL-treated mice are the same as those shown in Figure 7b.

**f, Pimonidazole alone does not elicit an appreciable increase in oxygen consumption in different experimental groups.** Wild-type and UCP1-KO mice acclimated to 30 °C (wild-type, n = 3; UCP1-KO, n = 4) or 4 °C (wild-type, n = 3; UCP1-KO, n = 4) were injected with pimonidazole alone. No appreciable increase in oxygen consumption was observed in any group. The wild-type mice acclimated to 4 °C are the same as those shown in panel e.

**g, CL induces a decrease in RER.** For clarity, only wild-type (filled circles) and UCP1-KO (open circles) mice acclimated to thermoneutrality are shown. These are the same mice as those shown in Figure 7a.

In panels e–g, each symbol represents one mouse. Values are means ± SEM.

**h–s, Representative confocal images of hypoxia adduct staining in IBAT from cold-acclimated wild-type, UCP1-KO, and UCP1$^{Myf5cKO}$ mice treated with CL followed by pimonidazole.** Tissues were stained for UCP1 (red; **h,l,p**), hypoxia adducts (green; **i,m,q**), and perilipin (blue; **j,n,r**). **k,o,s,** Merged images. CL treatment induced hypoxia adduct staining in IBAT from wild-type mice (panels h–k). In contrast, no hypoxia adduct staining was detected in UCP1-KO mice (panels l–o). In IBAT from UCP1$^{Myf5cKO}$ mice, only rare, small hypoxic regions were observed, confined close to residual UCP1-positive cells (panels p–s). Scale bar, 50 μm.

**Table S1.** Real-time qPCR primer sequences

| Gene | Forward (5' - 3') | Reverse (5' - 3') |
|---|---|---|
| *Ucp1* | GGCCTCTACGACTCAGTCCA | TAAGCCGGCTGAGATCTTGT |
| *Ckb* | AGTTCTCGGAGGTGCTCAAG | ATCAAACACCCCACCGACAG |
| *Alpl* | CCAACTCTTTTGTGCCAGAGA | GGCTACATTGGTGTTGAGCTTTT |
| *Atp2a2* | GCCACTCATGACAACCCACT | ACACAGCCGACGAAAGTCAG |
| *Ryr2* | CAAGGAAGGCTTGCTCCAGA | ACCTGGCAGTCACAGAGGTA |
| *Gk* | CCGCGAAGAAAGCAGTTCTG | CAAAAAACGTGTCGAGCTGGT |
| *Atgl* | TGACCATCTGCCTTCCAGA | TGTAGGTGGCGCAAGACA |
| *Dgat1* | TTATCGTGGTATCCTGAATTGGT | AAAAATAACCTTGCATTACTCAG |
| *Dgat2* | GGCGCTACTTCCGAGACTAC | TGGTCAGCAGGTTGTGTGTC |
| *Ant1* | GAGCTGCCTACTTCGGAGTC | CTGGGCAATCATCCAGCTCA |
| *Ant2* | GCTGGCCTGACTTCCTATCC | AAGCGTGCCTGTGTACATGA |
| *Pm20d1* | CCCCTGAACCCTCAGGACTT | ACTTCACCTGGTTCTGGTAGT |
| *TFIIB* | TGGAGATTTGTCCACCATGA | GAATTGCCAAACTCATCAAAACT |
| *18S rRNA* | AGTCCCTGCCCTTTGTACACA | CGATCCGAGGGCCTCACTA |
| *TFIIB* | TGGAGATTTGTCCACCATGA | GAATTGCCAAACTCATCAAAACT |
